# Towards understanding how we pay attention in naturalistic visual search settings

**DOI:** 10.1101/2020.07.30.229617

**Authors:** Nora Turoman, Ruxandra I. Tivadar, Chrysa Retsa, Micah M. Murray, Pawel J. Matusz

## Abstract

Research on attentional control has largely focused on single senses and the importance of behavioural goals in controlling attention. However, everyday situations are multisensory and contain regularities, both likely influencing attention. We investigated how visual attentional capture is simultaneously impacted by top-down goals, the multisensory nature of stimuli, *and* the contextual factors of stimuli’s semantic relationship and temporal predictability. Participants performed a multisensory version of the Folk et al. (1992) spatial cueing paradigm, searching for a target of a predefined colour (e.g. a red bar) within an array preceded by a distractor. We manipulated: 1) stimuli’s goal-relevance via distractor’s colour (matching vs. mismatching the target), 2) stimuli’s multisensory nature (colour distractors appearing alone vs. with tones), 3) the relationship between the distractor sound and colour (arbitrary vs. semantically congruent) and 4) the temporal predictability of distractor onset. Reaction-time spatial cueing served as a behavioural measure of attentional selection. We also recorded 129-channel event-related potentials (ERPs), analysing the distractor-elicited N2pc component both canonically and using a multivariate electrical neuroimaging framework. Behaviourally, arbitrary target-matching distractors captured attention more strongly than semantically congruent ones, with no evidence for context modulating multisensory enhancements of capture. Notably, electrical neuroimaging of surface-level EEG analyses revealed context-based influences on attention to both visual and multisensory distractors, in how strongly they activated the brain and type of activated brain networks. For both processes, the context-driven brain response modulations occurred long before the N2pc time-window, with topographic (network-based) modulations at ~30ms, followed by strength-based modulations at ~100ms post-distractor onset. Our results reveal that both stimulus meaning and predictability modulate attentional selection, and they interact while doing so. Meaning, in addition to temporal predictability, is thus a second source of contextual information facilitating goal-directed behaviour. More broadly, in everyday situations, attention is controlled by an interplay between one’s goals, stimuli’s perceptual salience, meaning and predictability. Our study calls for a revision of attentional control theories to account for the role of contextual and multisensory control.

## Introduction

Goal-directed behaviour depends on the ability to allocate processing resources towards the stimuli important to current behavioural goals (“attentional control”). On the one hand, our current knowledge about attentional control may be limited to the rigorous, yet artificial, conditions in which it is traditionally studied. On the other hand, findings from studies assessing attentional control with naturalistic stimuli (audiostories, films) may be limited by confounds from other processes present in such settings. Here, we systematically tested how traditionally studied goal- and salience-based attentional control interact with more naturalistic, context-based mechanisms.

In the real world, the location of goal-relevant information is rarely known in advance. Since the pioneering visual search paradigm (Treisman & Gelade, 1980), we know that in multi-stimulus settings target attributes can be used to control attention. Here, research provided conflicting results as to whether primacy in controlling attentional selection lies in task-relevance of objects’ attributes (Folk et al., 1992) or their bottom-up salience (e.g. Theeuwes, 1991). Folk et al. (1992) used a version of the spatial cueing paradigm and revealed that attentional capture is elicited only by distractors that matched the target colour. Consequently, they proposed the ‘task-set contingent attentional capture’ hypothesis, i.e., salient objects will capture attention only if they share features with the target and are thus potentially task-relevant. However, subsequently mechanisms beyond goal-relevance were shown to serve as additional sources of attentional control, e.g., spatiotemporal and semantic information within the stimulus and the environment where it appears (e.g., Chun & Jiang 1998; Peelen & Kastner, 2014; Summerfield et al., 2006; van Moorselaar & Slagter 2019; Press et al. 2020), and multisensory processes (Matusz & Eimer, 2011, 2013; Matusz et al. 2015a; Lunn et al. 2019; Soto-Faraco et al. 2019).

Some multisensory processes occur at early latencies (<100ms post-stimulus), generated within primary cortices (e.g., Talsma & Woldroff, 2005; Raij et al. 2010; Cappe et al. 2010; reviewed in de Meo et al., 2015; Murray et al. 2016a). This enables multisensory processes to influence attentional selection in a bottom-up fashion, potentially independently of the observer’s goals. This idea was supported by Matusz and Eimer (2011) who used a multisensory adaptation of Folk et al.’s (1992) task. The authors replicated the task-set contingent attentional capture effect and showed that visual distractors captured attention more strongly when accompanied by a sound, regardless of their goal-relevance. This demonstrated the importance of bottom-up multisensory enhancement for attentional selection of visual objects. However, interactions between such goals, multisensory influences on attentional control, and the stimuli’s temporal and semantic context^1^ remain unknown.

### Top-down contextual factors in attentional control

The temporal structure of the environment is routinely used by the brain to build predictions. Attentional control uses such predictions to improve the selection of target stimuli (e.g., Correa et al., 2005; Coull et al., 2000; Green & McDonald, 2010; Miniussi et al., 1999; Naccache et al., 2002; Rohenkohl et al., 2014; Tivadar et al. 2021) and the inhibition of task-irrelevant stimuli (here, location- and feature-based predictions have been more researched than temporal predictions; e.g., reviewed in Noonan et al. 2018; van Moorselaar & Slagter 2020a). In naturalistic, multisensory settings, temporal predictions are known to improve language comprehension (e.g. Luo & Poeppel, 2007; ten Oever & Sack, 2015), yet their role as a source of attentional control is less known (albeit see, Zion Golumbic et al. 2012, for their role in the “cocktail party” effect). Semantic relationships are another basic principle of organising information in real-world contexts. Compared to semantically incongruent or meaningless (arbitrary) multisensory stimuli, semantically congruent stimuli are more easily identified and remembered (e.g. Laurienti et al. 2004; Murray et al., 2004; Doehrmann & Naumer 2008; Chen & Spence, 2010; Matusz et al., 2015a; Tovar et al. 2020; reviewed in ten Oever et al. 2016; Murray et al., 2016b; Matusz et al. 2020) and also, more strongly attended (Matusz et al. 2015b, 2019a, 2019b; reviewed in Soto-Faraco et al., 2019; Matusz et al. 2019c). For example, Iordanenscu et al. (2009) demonstrated that search for naturalistic objects is faster when accompanied by irrelevant albeit congruent sounds.

What is unclear from existing research is the degree to which goal-based attentional control interacts with salience-driven (multisensory) mechanisms *and* such contextual factors. Researchers have been clarifying such interactions, but typically in a pair-wise fashion, between e.g., attention and semantic memory, or attention and predictions (reviewed in Summerfield & Egner 2009; Nobre & Gazzaley 2016; Press et al. 2020). However, in everyday situations these processes do not interact in an orthogonal, but, rather, a synergistic fashion, with multiple sources of control interacting simultaneously (ten Oever et al. 2016; Nastase et al. 2020). Additionally, in the real world, these processes operate on both unisensory and multisensory stimuli, where the latter are often more perceptually salient than the former (e.g., Santangelo & Spence 2007; Matusz & Eimer 2011). Thus, one way to create more complete and “naturalistic” theories of attentional control is by investigating how one’s goals interact with *multiple* contextual factors in controlling attentional selection – and doing so in *multisensory settings*.

### The present study

To shed light on how attentional control operates in naturalistic visual search settings, we investigated how visual and multisensory attentional control interact with distractor temporal predictability and multisensory semantic relationship when all are manipulated simultaneously. We likewise set out to identify brain mechanisms supporting such complex interactions. To address these questions in a rigorous and state-of-the-art fashion, we employed a ‘naturalistic laboratory’ approach that builds on several methodological advances (Matusz et al., 2019c). First, we used a paradigm that isolates a specific cognitive process, i.e., Matusz and Elmer’s (2011) multisensory adaptation of the Folk et al.’s (1992) task, where we additionally manipulated distractors’ temporal predictability and relationship between their auditory and visual features. In Folk et al.’s task, attentional control is measured via well-understood spatial cueing effects, where larger effects (e.g., for target-colour and audiovisual distractors) reflect stronger attentional capture. Notably, distractor-related responses have the added value as they isolate attentional from later, motor response-related, processes. Second, we measured a well-researched brain correlate of attentional object selection, the N2pc event-related potential (ERP) component. The N2pc is a negative-going voltage deflection starting at around 200ms post-stimulus onset at posterior electrode sites contralateral to stimulus location (Luck & Hillyard, 1994a, 1994b; Eimer, 1996; Girelli & Luck, 1997). Studies canonically analysing N2pc have provided strong evidence for task-set contingence of attentional capture (e.g., Kiss et al., 2008a; 2008b; Eimer et al., 2009). Importantly, N2pc is also sensitive to meaning (e.g., Wu et al., 2015) and predictions (e.g., Burra & Kerzel, 2013), whereas its sensitivity to multisensory enhancement is limited (van der Burg et al. 2011, but see below). This joint evidence makes the N2pc a valuable ‘starting point’ for investigating interactions between visual goals and more naturalistic sources of control. Third, analysing the traditional EEG markers of attention with advanced frameworks like electrical neuroimaging (e.g., Lehmann & Skrandies 1980; Murray et al., 2008; Tivadar & Murray 2019) might offer an especially robust, accurate and informative approach.

Briefly, an electrical neuroimaging framework encompasses multivariate, reference-independent analyses of global features of the electric scalp field. Its main added value is that it readily distinguishes the neurophysiological mechanisms driving differences in ERPs across experimental conditions in *surface-level* EEG: 1) “gain control” mechanisms, modulating the strength of activation within an indistinguishable brain network, and 2) topographic (network-based) mechanisms, modulating the recruited brain sources (scalp EEG topography differences forcibly follow from changes in the underlying sources; Murray et al. 2008). Electrical neuroimaging overcomes interpretational limitations of canonical N2pc analyses. Most notably, a difference in mean N2pc amplitude can arise from both strength-based and **topographic** mechanisms (albeit it is assumed to signify gain control); it can also emerge from different brain source configurations (for a full discussion, see Matusz et al., 2019b).

We recently used this approach to better understand brain and cognitive mechanisms of attentional control. We revealed that distinct brain networks are active during ~N2pc time-window during visual goal-based *and* multisensory bottom-up attention control (across the lifespan; Turoman et al. 2021a, 2021b). However, these reflect spatially-selective, lateralised brain mechanisms, partly captured by the N2pc (via the contra- and ipsilateral comparison). There is little existing evidence to strongly predict how interactions between goals, stimulus salience and context can occur in the brain. Schröger et al. (2015) proposed that temporally unpredictable events attract attention more strongly (to serve as a signal to reconfigure the predictive model about the world), visible in larger behavioural responses and ERP amplitudes. Both predictions and semantic memory could be used to reduce attention to known (i.e., less informative) stimuli. Indeed, goal-based control uses knowledge to facilitate visual and multisensory processing (Summerfield et al. 2008; Iordanescu et al., 2008; Matusz et al. 2016; Sarmiento et al. 2016). However, several questions remain. Does knowledge affect attention to task-*irrelevant* stimuli the same way? How early do contextual factors influence stimulus processing here, if both processes are known to do so <150ms post-stimulus (Summerfield & Egner, 2009; ten Oever et al. 2016). Finally, do contextual processes operate through lateralised or non-lateralised brain mechanisms? Below we specify our hypotheses.

We expected to replicate the TAC^2^ effect: In behaviour, visible as large behavioural capture for target-colour matching distractors and no capture for nontarget-colour matching distractors (e.g., Folk et al., 1992; Folk, et al., 2002; Lien et al., 2008); in canonical EEG analyses - enhanced N2pc amplitudes for target-colour than nontarget-colour distractors (Eimer et al., 2009). TAC should be modulated by both contextual factors: the predictability of distractor onset and the multisensory relationship between distractor features (semantic congruence vs. arbitrary pairing; Wu et al. 2015; Burra & Kerzel, 2013). However, as discussed above, we had no strong predictions how the contextual factors would modulate TAC (or if they interact while doing so), as these effects have never been tested systematically together, on audio-visual and task-irrelevant stimuli. For multisensory enhancement of capture, we expected to replicate it behaviourally (Matusz & Eimer 2011), but without strong predictions about concomitant N2pc modulations (c.f. van der Burg et al. 2011). We expected multisensory enhancement of capture to be modulated by contextual factors, especially multisensory relationship, based on the extensive literature on the role of semantic congruence in multisensory cognition (Doehrmann & Naumer, 2008; ten Oever et al. 2016). Again, we had no strong predictions as to the directionality of these modulations or interaction of their influences.

We were primarily interested if interactions between visual goals (task-set contingent attentional capture, TAC), multisensory salience (multisensory enhancement of capture, MSE) and contextual processes are supported by strength-based (i.e., “gain”-like; i.e., one network is active more strongly for some and less strongly for other experimental conditions) and/or topographic (i.e., different networks are activated for different experimental conditions) brain mechanisms, as observable in *surface-level* EEG data when using multivariate analyses like electrical neuroimaging. The second aim of our study was to clarify if the attentional and contextual control interactions are supported by lateralised (N2pc-like) or nonlateralized mechanisms. To this aim, we analysed if those interactions are captured by canonical N2pc analyses or electrical neuroimaging analyses of the lateralised distractor-elicited ERPs ~180-300ms post-stimulus (N2pc-like time-window). These analyses would reveal presence of strength- and topographic *spatially-selective* brain mechanisms contributing to attentional control. However, analyses of the N2pc assume not only lateralised activity, but also symmetry; in brain anatomy but also in scalp electrodes, detecting homologous brain activity over both hemispheres. This may prevent them from detecting other, less-strongly-lateralised brain mechanisms of attentional control. We have previously found nonlateralised mechanisms to play a role in attentional control in multisensory settings (Matusz et al. 2019b). Also, semantic information and temporal expectations (and feature-based attention) are known to modulate nonlateralised ERPs (Saenz et al. 2003; Dell’Acqua et al. 2010; Dassanayake et al. 2016). Thus, as the third aim of our study, we investigated whether contextual control affects stages associated with attentional selection (reflected by the N2pc) or also earlier processing stages. We tested this by measuring strength- and/or topographic nonlateralised brain mechanisms across the whole post-stimulus time-period activity.

## Materials and Methods

### Participants

Thirty-nine adult volunteers participated in the study (5 left-handed, 14 males, *M_age_*: 27.5 years, *SD*: 4 years, range: 22–38 years). We conducted post-hoc power analyses for the two effects that have been previously behaviourally studied with the present paradigm, namely TAC and MSE. Based on the effect sizes in the original Matusz and Eimer (2011, Exp.2), the analyses revealed sufficient statistical power for both behavioural effects with the collected sample. For ERP analyses, we could calculate power analyses only for the TAC effect. Based on a purely visual ERP study (Eimer et al., 2009) we revealed there to be sufficient statistical power to detect TAC in the N2pc in the current study (all power calculations are available in the Supplemental Online Materials, SOMs). Participants had normal or corrected-to-normal vision and normal hearing and reported no prior or current neurological or psychiatric disorders. Participants provided informed consent before the start of the testing session. All research procedures were approved by the Cantonal Commission for the Ethics of Human Research (CER-VD; no. 2018-00241).

### Task properties and procedures

#### General task procedures

The full experimental session consisted of participants completing four experimental Tasks. All the Tasks were close adaptations of the original paradigm of Matusz and Eimer (2011 Exp.2; that is, in turn, an adaptation of the spatial-cueing task of Folk et al. [1992]). Across all the Tasks, the instructions and the overall experimental set up were the same as in the study of Matusz & Eimer (1992, Exp.2; see Figure 1A). Namely, participants searched for a target of a predefined colour (e.g., a red bar) in a 4-element array, and assessed the target’s orientation (vertical vs. horizontal). Furthermore, in all Tasks, the search array was always preceded by an array containing colour distractors. Those distractors always either matched the target colour (red set of dots) or matched another, nontarget colour (blue set of dots); on 50% of all trials the colour distractors would be accompanied by a sound (audiovisual distractor condition). The distractor appeared in each of the four stimulus locations with equal probability (25%) and was thus not predictive of the location of the incoming target. Differences in response speed on trials where distractor and target appeared in the same vs. different locations were used to calculate behavioural cueing effects that were the basis of our analyses (see below). Like in the Matusz and Eimer (2011) study, across all Tasks, each trial consisted of the following sequence of arrays: base array (duration manipulated; see below), followed by distractor array (50ms duration), followed by a fixation point (150ms duration), and finally the target array (50ms duration, see Figure 1A).

**Figure 1.**
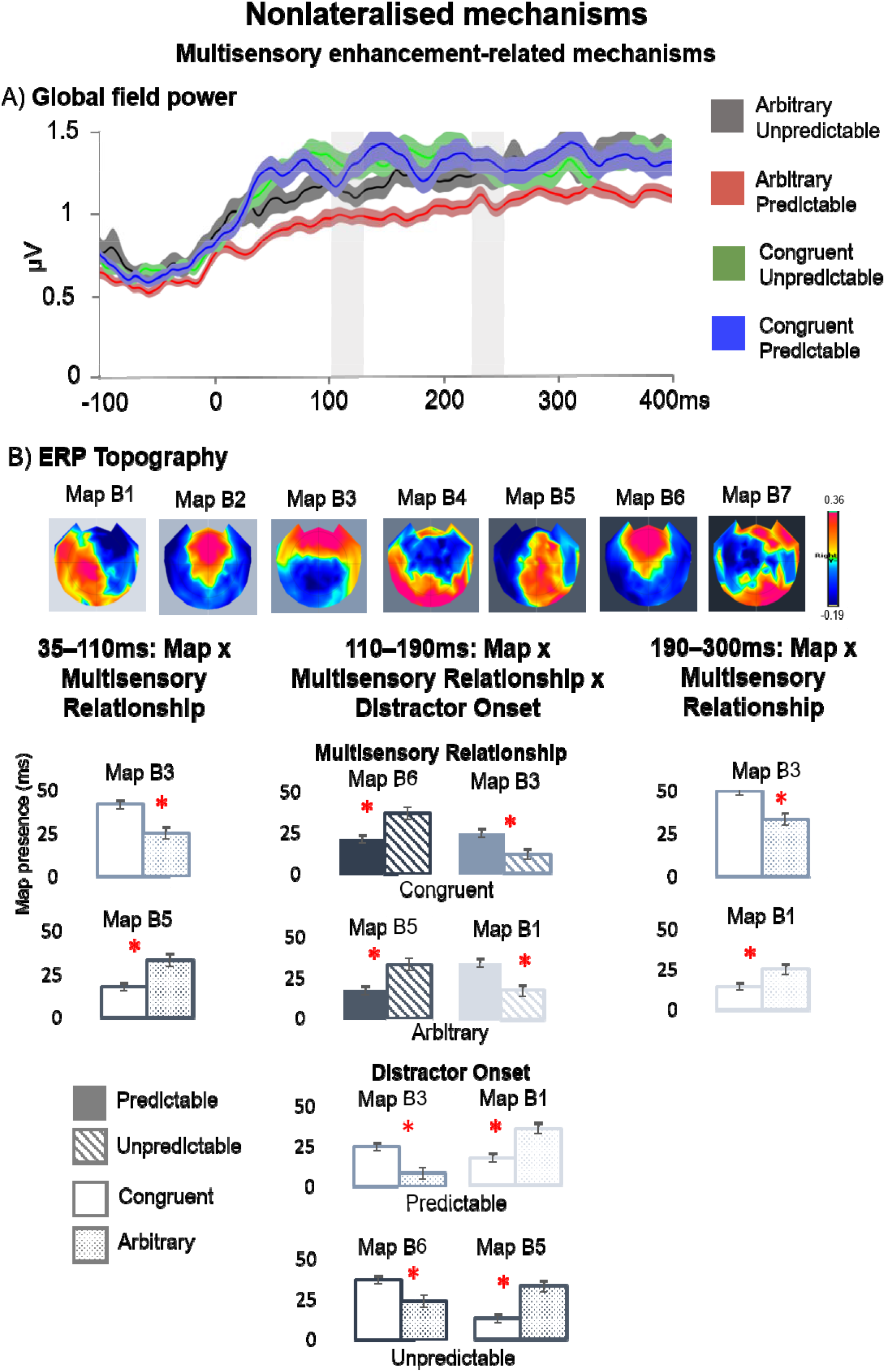
**A)** An example trial of the general experimental “Task” is shown, with four successive arrays. The white circle around the target location (here the target is a blue diamond) and the corresponding distractor location serves to highlight, in this case, a target-matching distractor colour condition, with a concomitant sound, i.e., TCCAV. **B)** The order of Tasks, with the corresponding conditions of Multisensory Relationship in red, and Distractor Onset in green, shown separately for each Task, in the successive order in which they appeared in the study. Under each condition, its operationalisation is given in brackets in the corresponding colour. Predictable and unpredictable blocks before and after the training (1 & 2 and 3 & 4, respectively) were counterbalanced across participants. **C)** Events that were part of the Training. Association phase: an example pairing option (red – high pitch, blue – low pitch) with trial progression is shown. Testing phase: the pairing learnt in the Association phase would be tested using a colour word or a string of x’s in the respective colour. Participants had to indicate whether the pairing was correct via a button press, after which feedback was given.

The differences to the original study involved the changes necessary to implement the two new, contextual factors that were manipulated across the four Tasks (Figure 1B).^3^ To implement the *Multisensory Relationship* factor, after the first two Tasks, participants completed a training session (henceforth *Training*), after which they completed the remaining two Tasks. To implement the *Distractor Onset* factor, the predictability of the onset of the distractors was manipulated, being either stable (as in the original study, Tasks 2 and 4) or varying between three durations (Tasks 1 and 3). The setup involving 4 consecutive Tasks separated by Training allowed a systematic comparison between the four levels of the two contextual factors. We now describe in more detail the procedures related to all Tasks, after which we provide more details on the different tasks themselves.

The base array contained four differently coloured sets of closely aligned dots, each dot subtending 0.1° × 0.1° of visual angle. The sets of dots were spread equidistally along the circumference of an imaginary circle against a black background, at an angular distance of 2.1° from a central fixation point. Each set could be of one of four possible colours (according to the RGB scale): green (0/179/0), pink (168/51/166), gold (150/134/10), silver (136/136/132). In the distractor array, one of the base array elements changed colour to either a target-matching colour, or a target-nonmatching colour that was not present in any of the elements before. The remaining three distractor array elements did not change their colour. The distractors and the subsequent target diamonds could have either a blue (RGB values: 31/118/220) or red (RGB values: 224/71/52) colour. The target array contained four bars (rectangles), where one was always the colour-defined target. The target colour was counterbalanced across participants. Target orientation (horizontal or vertical) was randomly determined on each trial. The two distractor colours were randomly selected with equal probability before each trial, and the location of the colour change distractor was not spatially predictive of the subsequent target location (distractor and target location were the same on 25% of trials). On half of all trials, distractor onset coincided with the onset of a pure sine-wave tone, presented from two loudspeakers on the left and right sides of the monitor. Sound intensity was 80 dB SPL (as in Matusz & Eimer, 2011), measured using an audiometer placed at a position adjacent to participants’ ears (CESVA SC160). Through manipulations of the in-/congruence between distractor and target colour and of the presence/absence of sound during distractor presentations, there were four types of distractors, across all the Tasks: visual distractors that matched the target colour (TCCV, short for *target-colour cue visual*), visual distractors that did not match the target colour (NCCV, *nontarget-colour cue visual*), audiovisual distractors that matched the target colour (TCCAV, *target-colour cue audiovisual*), and audiovisual distractors that did not match the target colour (NCCAV, *nontarget-colour cue, audiovisual*).

The experimental session consisted of 4 Tasks, each spanning 8 blocks of 64 trials. This resulted in 2,048 trials in total (512 trials per Task). Participants were told to respond as quickly and accurately as possible to the targets’ orientation by pressing one of two horizontally aligned round buttons (Lib Switch, Liberator Ltd.) that were fixed onto a tray bag on the participants’ lap. If participants did not respond within 5000ms of the target onset, next trial was initiated; otherwise the next trial was initiated immediately after the button press. Feedback on accuracy was given after each block, followed by a progress screen (*a treasure map*), which informed participants of the number of remaining blocks and during which participants could take a break. Breaks were also taken between each Task, and before and after the Training. As a pilot study revealed sufficient proficiency at conducting the tasks after a few trials (over 50% accuracy), participants did not practice doing the Tasks before administration unless they had trouble following the task instructions. The experimental session took place in a dimly lit, sound-attenuated room, with participants seated at 90cm from a 23” LCD monitor with a resolution of 1080 × 1024 (60-Hz refresh rate, HP EliteDisplay E232). All visual elements were approximately equiluminant (~20cd/m^2^), as determined by a luxmeter placed at a position close to the screen, measuring the luminance of the screen filled with each respective element’s colour. The averages of three measurement values per colour were averaged across colours and transformed from lux to cd/m^2^ to facilitate comparison with the results of Matusz & Eimer (2011). The experimental session lasted <3h in total, including an initial explanation and obtaining consent, EEG setup, administration of Tasks and Training, and breaks.

We now describe the details of the Tasks and Training, which occurred always in the same general order: Tasks 1 and 2, followed by the Training, followed by Tasks 3 and 4 (the order of Tasks 1 and 2 and, separately, the order of Tasks 3 and 4, was counterbalanced across participants). Differences across the four Tasks served to manipulate the two contextual factors (illustrated in Figure 1B). The factor *Multisensory Relationship* represented the relation between the visual (the colour of the distractor) and the auditory (the accompanying sound) component stimuli that made up the distractors. These two stimuli could be related just by their simultaneous presentation (Arbitrary condition) or by additionally sharing meaning (Congruent condition). The factor *Distractor Onset* represented the temporal predictability of the distractors, i.e., whether their onset was constant within Tasks and, therefore Predictable condition, or variable and, therefore, Unpredictable condition. The manipulation of the two context factors led to the creation of four contexts, represented by each of the Tasks 1–4 (i.e., Arbitrary Unpredictable, Arbitrary Predictable, Congruent Unpredictable, and Congruent Predictable). To summarise, the two within-task factors encompassing distractor colour and tone presence/absence, together with the two between-task factors resulted in a total of four factors in our analysis design: Distractor Colour (TCC vs. NCC), Distractor Modality (V vs. AV), Distractor Onset (Predictable vs. Unpredictable) and Multisensory Relationship (Arbitrary vs. Congruent)^4^.

#### Tasks 1 and 2

As mentioned above, across Tasks 1 and 2, the colour of the distractor and the sound accompanying the colour distractor were related only by their simultaneous presentation. As such, trials from Tasks 1 and 2 made up the Arbitrary condition of the Multisensory Relationship factor. Sound frequency was always 2000Hz (as in Matusz & Eimer, 2011). The main difference between Task 1 and Task 2 lied in the onset of the distractors in those tasks. Unbeknownst to participants, in Task 1, duration of the base array varied randomly on a trial-by-trial basis, between 100ms, 250ms and 450ms, i.e., the distractor onset was unpredictable. In contrast, in Task 2, the base array duration was always constant, at 450ms, i.e., the distractor onset was predictable. With this manipulation, considering the between-task factors: Task 1 represented Arbitrary (Multisensory Relationship) and Unpredictable (Distractor Onset) trials, and Task 2 - Arbitrary (Multisensory Relationship) and Predictable (Distractor Onset) trials.

#### Training

The Training served to induce in participants a semantic-level association between a specific distractor colour and a specific pitch. This rendered distractors in the Tasks following the Training semantically related (Congruent), and distractors in the preceding Tasks semantically unrelated (Arbitrary). The Training consisted of an Association phase followed by a Testing phase (both based on the association task in Sui, He & Humphreys, 2012; see also Sun et al., 2016).

##### I. Association phase

The Association phase served to induce the AV associations in participants. Participants were shown alternating colour word–pitch pairs, presented in the centre of the screen (the tone was presented from two lateral speakers, rendering it spatially diffuse and so appearing to also come from the centre of the screen). The words denoted one of two distractor colours (*red* or *blue*). The tone of either high (4000Hz) or low (300Hz) pitch. Both the colour word and sound were presented for 2 seconds, after which a central fixation cross was presented for 150ms, followed by the next colour word–pitch pair. There could be two possible colour–pitch pairing options. In one, the high-pitch tone was associated with the word *red*, the low-pitch tone - with the word *blue*. In the second option, the high-pitch tone was associated with the word *blue*, the low-pitch tone with the word *red* (see Figure 1C, Association phase). Pairing options were counterbalanced across participants. Thus, for participants trained with the first option, the pairing of word *red* and a high-pitch tone would be followed by the pairing of the word *blue* with a low-pitch tone, again followed by the red–high pitch pairing, etc. There were 10 presentations per pair, resulting in a total of 20 trials. Colour words were chosen instead of actual colours to ensure that the AV associations were based on meaning rather than lower-level stimulus features (for examples of such taught crossmodal correspondences see, e.g., Ernst, 2007). Also, colour words were shown in participants’ native language (speakers: 19 French, 8 Italian, 5 German, 4 Spanish, 3 English). Participants were instructed to try to memorise the pairings as best as they could, being informed that they would be subsequently tested on how well they learnt the pairings.

##### II. Testing phase

The Testing phase served to ensure that the induced colour–pitch associations was strong. Now, participants were shown colour word–pitch pairings (as in the Association phase) but also colour–pitch pairings (a string of x’s in either red or blue, paired with a sound, Figure 1C, *Testing phase* panel). Additionally, now, the pairings either matched or mismatched the type of associations induced in the Association phase, e.g., if the word *red* have been paired with a high-pitch tone in the Testing phase, the matching pair now would be a word *red* or red x’s, paired with a high-pitch tone, and mismatching pair - the word *red* or red x’s paired with a low-pitch tone. Participants had to indicate if a given pair was matched or mismatched by pressing one of two buttons (same button setup as in the Tasks). Participants whose accuracy was ≤50% had to repeat the testing.

The paradigm that Sui et al. (2012) have designed led to people being able to reliably associate low-level visual features (colours, geometric shapes) with abstract social concepts (themselves, their friend, a stranger). Following their design, in the Testing phase, each pairing was shown for 250ms, of which 50ms was the sound (instead of the stimulus duration of 100ms that Sui et al. used, to fit our stimulus parameters). The pairing presentation was followed by a blank screen (800ms), during which participants had to respond, and after each responses a screen with feedback on their performance appeared. Before each trial, a fixation cross was also shown, for 500ms. Each participant performed three blocks of 80 trials, with 60 trials per possible combination (colour word – sound matching, colour word – sound mismatching, colour – sound matching, colour – sound mismatching). A final summary of correct, incorrect, and missed trials was shown at the end of Testing phase.

#### Tasks 3 and 4

Following the Training, in Tasks 3 and 4, the distractors’ colour and the accompanying sound were now semantically related. Thus, the trials from these two Tasks made up the (semantically) Congruent condition of the Multisensory Relationship factor. Only congruent colour–pitch distractor pairings were now presented, as per the pairing option induced in the participants. That is, if the colour red was paired with a high-pitch tone in the Association phase, red AV distractors in Tasks 3 and 4 were always accompanied by a high-pitch tone. The pitch of sounds was now either 300Hz (low-pitch condition; chosen based on Matusz & Eimer, 2013, where two distinct sounds were used) or 4000Hz (high-pitch condition; chosen for its comparable perceived loudness in relation to the above two sound frequencies, as per the revised ISO 226:2003 equal-loudness-level contours standard; Spierer et al. 2013). As between Tasks 1 and 2, Task 3 and Task 4 differed in the predictability of distractor onsets, i.e., in Task 3, distractor onset was unpredictable, and in Task 4 - predictable. Therefore, Task 3 represented Congruent (Multisensory Relationship) and Unpredictable (Distractor Onset) trials, and Task 4 - Congruent (Multisensory Relationship) and Predictable (Distractor Onset) trials.

### EEG acquisition and preprocessing

Continuous EEG data sampled at 1000Hz was recorded using a 129-channel HydroCel Geodesic Sensor Net connected to a NetStation amplifier (Net Amps 400; Electrical Geodesics Inc., Eugene, OR, USA). Electrode impedances were kept below 50kΩ, and electrodes were referenced online to Cz. First, offline filtering involved a 0.1Hz high-pass and 40Hz low-pass as well as 50Hz notch (all filters were second-order Butterworth filters with −12dB/octave roll-off, computed linearly with forward and backward passes to eliminate phase-shift). Next, the EEG was segmented into peri-stimulus epochs from 100ms before distractor onset to 500ms after distractor onset. An automatic artefact rejection criterion of ±100μV was used, along with visual inspection. Epochs were then screened for transient noise, eye movements, and muscle artefacts using a semi-automated artefact rejection procedure. Data from artefact contaminated electrodes were interpolated using three-dimensional splines (Perrin et al., 1987). Across all Task, 11% of epochs were removed on average and 8 electrodes were interpolated per participant (6% of the total electrode montage).

Cleaned epochs were averaged, baseline corrected to the 100ms pre-distractor time interval, and re-referenced to the average reference. Next, to eliminate residual environmental noise in the data, a 50Hz filter was applied^5^. All the above steps were done separately for ERPs from the four distractor conditions, and separately for distractors in the left and right hemifield. We next relabeled ERPs from certain conditions, as is done in traditional lateralised ERP analyses (like those of the N2pc). Namely, we relabelled single-trial data from all conditions where distractors appeared on the *left* so that the electrodes over the left hemiscalp now represented the activity over the right hemiscalp, and electrodes over the right hemiscalp – represented activity over the left hemiscalp, thus creating “mirror distractor-on-the-right” single-trial data. Next, these mirrored data and the veridical “distractor-on-the-right” data from each of the 4 distractor conditions were averaged together, creating a single average ERP for each of the 4 distractor conditions. The contralaterality factor (i.e. contralateral vs. ipsilateral potentials) is normally represented by separate ERPs (one for contralateral activity, and one for ipsilateral activity; logically more pairs for pair-wise N2pc analyses). In our procedure, the lateralised voltage gradients across the whole scalp are preserved within each averaged ERP by simultaneous inclusion of both contralateral and ipsilateral hemiscalp activation. Such a procedure enabled us to fully utilise the capability of the electrical neuroimaging analyses in revealing both lateralised and non-lateralised mechanisms that support the interactions of attentional control with context control. As a result of the relabelling, we obtained 4 different ERPs: TCCV (target colour-cue, Visual), NCCV (nontarget colour-cue, Visual), TCCAV (target colour-cue, AudioVisual), NCCAV (nontarget colour-cue, Audiovisual). Preprocessing and EEG analyses, unless otherwise stated, were conducted using CarTool software (available for free at www.fbmlab.com/cartool-software/; Brunet, Murray, & Michel, 2011).

### Data analysis design

#### Behavioural analyses

Like in Matusz and Eimer (2011), and because mean reaction times (RTs) and accuracy did not differ significantly between the four Tasks, the basis of our analyses was RT spatial cueing effects (henceforth “behavioural capture effects”). These were calculated by subtracting the mean RTs for trials where the distractor and target were in the same location from the mean RTs for trials where the distractor and the target location differed, separately for each of the four distractor conditions. Such spatial cueing data were analysed using the repeated-measures analysis of variance (rmANOVA). Error rates (%) were also analysed. As they were not normally distributed, we analysed error rates using the Kruskal–Wallis *H* test and the Durbin test. The former was used to analyse if error rates differed significantly between Tasks, while the latter was used to analyse differences between experimental conditions within each Task separately.

Following Matusz and Eimer (2011), RT data were cleaned by discarding incorrect and missed trials, as well as RTs below 200ms and above 1000ms. Additionally, to enable more direct comparisons with the developmental study for which current Task 1 served as an adult control (Turoman et al., 2021a, 2021b), we have further removed trials with RTs outside 2.5SD of the individual mean RT. As a result, a total of 5% of trials across all Tasks were removed. Next, behavioural capture effects were submitted to a four-way 2 × 2 × 2 × 2 rmANOVA with factors: Distractor Colour (TCC vs. NCC), Distractor Modality (V vs. AV), Multisensory Relationship (Multisensory Relationship; Arbitrary vs. Congruent), and Distractor Onset (Distractor Onset; Unpredictable vs. Predictable). Due to the error data not fulfilling criteria for normality, we used Distractor-Target location as a factor in the analysis, conducting 3-way Durbin tests for each Task, with factors Distractor Colour, Distractor Modality, and Distractor-Target Location. All analyses, including post-hoc paired *t*-tests, were conducted using SPSS for Macintosh 26.0 (Armonk, New York: IBM Corporation). For brevity, we only present the RT results in the Results, and the error rate results can be found in SOMs.

#### ERP analyses

The preprocessing of the ERPs triggered by the visual and audiovisual distractors across the 4 different experimental blocks created ERP averages in which the contralateral versus ipsilateral ERP voltage gradients across the whole scalp were preserved. We first conducted a canonical N2pc analysis, as the N2pc is a well-studied and well-understood correlate of attentional selection in visual settings. However, it is unclear if the N2pc also indexes bottom-up attentional selection modulations by multisensory stimuli, or top-down modulations by contextual factors like multisensory semantic relationships (for visual-only study, see e.g., Wu et al. 2015) or stimulus onset predictability (for visual-only study, see e.g., Burra & Kerzel, 2013). N2pc analyses served also to bridge electrical neuroimaging analyses with the existing literature and EEG approaches more commonly used to investigate attentional control. Briefly, electrical neuroimaging encompasses a set of multivariate, reference-independent analyses of global features of the electric field measured at the scalp (König et al., 2014; Michel & Murray, 2012; Murray, Brunet, & Michel, 2008; Lehmann & Skrandies, 1980; Tivadar & Murray, 2019; Tzovara et al., 2012) that can detect spatiotemporal patterns in EEG across different contexts and populations (e.g., Neel et al. 2019; Matusz et al. 2018). The key advantages of electrical neuroimaging analyses over canonical N2pc analyses and how the former can complement the latter when combined, are described in the Introduction.

##### Canonical N2pc analysis

To analyse lateralised mechanisms using the traditional N2pc approach, we extracted mean amplitude values from, first, two electrode clusters comprising PO7/8 electrode equivalents (e65/90; most frequent electrode pair used to analyse the N2pc), and, second, their six immediate surrounding neighbours (e58/e96, e59/e91, e64/e95, e66/e84, e69/e89, e70/e83), over the 180–300ms post-distractor time-window (based on time-windows commonly used in traditional N2pc studies, e.g., Luck & Hillyard, 1994b; Eimer, 1996; including distractor-locked N2pc, Eimer & Kiss 2008; Eimer et al. 2009). Analyses were conducted on the mean amplitude of the N2pc difference waveforms, which were obtained by subtracting the average of amplitudes in the ipsilateral posterior-occipital cluster from the average of amplitudes in the contralateral posterior-occipital cluster. This step helped mitigate the loss of statistical power that could result from the addition of contextual factors into the design. N2pc means were thus submitted to a 4-way 2 × 2 × 2 × 2 rmANOVA with factors Distractor Colour (TCC vs. NCC), Distractor Modality (V vs. AV), Multisensory Relationship (Arbitrary vs. Congruent), and Distractor Onset (Unpredictable vs. Predictable), analogously to the behavioural analysis. Notably, the N2pc is not sensitive to the location of the stimulus of interest *per se*, but rather to the side of its presentation. As such, in canonical analyses of distractor-elicited N2pc, the congruence between distractor and target, unlike in behavioural analyses, is not considered (e.g., Lien et al. 2008; Eimer & Kiss 2008; Eimer et al. 2009). Consequently, in our N2pc analyses, target-location congruent and incongruent distractor ERPs were averaged, as a function of the side of distractor presentation.

##### Electrical Neuroimaging of the N2pc component

Our electrical neuroimaging analyses separately tested response strength and topography in N2pc-like lateralised ERPs (see e.g. Matusz et al., 2019b for a detailed, tutorial-like description of how electrical neuroimaging measures can aid the study of attentional control processes). We assessed if interactions between visual goals, multisensory salience and contextual factors 1) modulated the distractor-elicited lateralised ERPs, and 2) if they do so by altering the strength of responses within statistically indistinguishable brain networks and/or altering the recruited brain networks.

##### I. Lateralised analyses

To test for the involvement of strength-based spatially-selective mechanisms, we analysed Global Field Power (GFP) in lateralised ERPs. GFP is the root mean square of potential [μV] across the entire electrode montage (see Lehmann & Skrandies, 1980). To test for the involvement of network-related spatially-selective mechanisms, we analysed stable patterns in ERP topography characterising different experimental conditions using a clustering approach known as the Topographic Atomize and Agglomerate Hierarchical Clustering (TAAHC). This topographic clustering procedure generated sets of clusters of topographical maps that explained certain amounts of variance within the group-averaged ERP data. Each cluster was labelled with a ‘template map’ that represented the centroid of its cluster. The optimal number of clusters is one that explains the largest global explained variance in the group-averaged ERP data with the smallest number of template maps, and which we identified using the modified Krzanowski–Lai criterion (Murray et al., 2008). In the next step, i.e., the so-called fitting procedure, the single-subject data was ‘fitted’ back onto the topographic clustering results, such that each datapoint of each subject’s ERP data over a chosen time-window was labelled by the template map with which it was best spatially correlated. This procedure resulted in a number of timeframes that a given template map was present over a given time-window, which durations (in milliseconds) we then submitted to statistical analyses described below.

In the present study, we conducted strength- and topographic analyses using the same 4-way repeated-measures design as in the behavioural and canonical N2pc analyses, on the lateralised whole-montage ERP data. Since the N2pc is a lateralised ERP, we first conducted an electrical neuroimaging analysis of lateralised ERPs in order to uncover the modulations of the N2pc by contextual factors. To obtain *global* electrical neuroimaging measures of *lateralised* N2pc effects, we computed a difference ERP by subtracting the voltages over the contralateral and ipsilateral hemiscalp, separately for each of the 4 distractor conditions. This resulted in a 59-channel difference ERP (as the midline electrodes from the 129-electrode montage were not informative). Next, this difference ERP was mirrored onto the other side of the scalp, recreating a “fake” 129 montage (with values on midline electrodes now set to 0). It was on these mirrored “fake” 129-channel lateralised difference ERPs that lateralised strength-based and topography-based electrical neuroimaging analyses were performed. Here, GFP was extracted over the canonical 180–300ms N2pc time-window and submitted to a 2×2×2×2 rmANOVA with factors Distractor Colour (TCC vs. NCC), Distractor Modality (V vs. AV), as well as the two new factors, Multisensory Relationship (Arbitrary vs. Congruent), and Distractor Onset (Distractor Onset; Unpredictable vs. Predictable). Meanwhile, for topographic analyses, the “fake” 129-channel data across the 4 Tasks were submitted to a topographic clustering over the entire post-distractor period. Next, the data were fitted back over the 180-300ms period. Finally, the resulting number of timeframes (in ms) was submitted to the same rmANOVA as the GFP data above.

It remains unknown if the tested contextual factors modulate lateralised ERP mechanisms at all. Given evidence that semantic information and temporal expectations can modulate *nonlateralised* ERPs within the first 100–150ms post-stimulus (e.g., Dell’Acqua et al., 2010; Dassanayake et al., 2016), we also investigated the influence of contextual factors on nonlateralised voltage gradients, in an exploratory fashion. It must be noted that ERPs are sensitive to the inherent physical differences in visual and audiovisual conditions. Specifically, on audiovisual trials, the distractor-induced ERPs would be contaminated by brain response modulations induced by sound processing, with these modulations visible in our data already at 40ms post-distractor. Consequently, any direct comparison of visual-only and audiovisual ERPs would index auditory processing per se and not capture of attention by audiovisual stimuli. Such confounded sound-related activity is eliminated in the canonical N2pc analyses through the contralateral-minus-ipsilateral subtraction. To eliminate this confound in our electrical neuroimaging analyses here, we calculated difference ERPs, first between TCCV and NCCV conditions, and then between TCCAV and NCCAV conditions. Such difference ERPs, just as the canonical N2pc difference waveform, subtract out the sound processing confound in visually-induced ERPs. As a result of those difference ERPs, we removed factors Distractor Colour and Distractor Modality, and produced a new factor, Target Difference (two levels: D_AV_ [TCCAV – NCCAV difference] and D_v_ [TCCV – NCCV difference]), that indexed the enhancement of visual attentional control by sound presence.

##### II. Nonlateralised analyses

All nonlateralised electrical neuroimaging analyses involving context factors were based on the Target Difference ERPs. Strength-based analyses, voltage and GFP data were submitted to 3-way rmANOVAs with factors: Multisensory Relationship (Arbitrary vs. Congruent), Distractor Onset (Unpredictable vs. Predictable), and Target Difference (D_AV_ vs. D_V_), and analysed using the STEN toolbox 1.0 (available for free at https://zenodo.org/record/1167723#.XS3lsi17E6h). Follow-up tests involved further ANOVAs and pairwise *t*-tests. To correct for temporal and spatial correlation (see Guthrie & Buchwald, 1991), we applied a temporal criterion of >15 contiguous timeframes, and a spatial criterion of >10% of the 129-channel electrode montage at a given latency for the detection of statistically significant effects at an alpha level of 0.05. As part of topography-based analyses, we segmented the ERP difference data across the post-distractor and pre-target onset period (0 – 300ms from distractor onset). To isolate the effects related to each of the two cognitive processes and reduce the complexity of the performed analyses, we carried out two topographic clustering analyses. Topographic clustering on nonlinear mechanisms contributing to TAC was based on the visual Target Difference ERPs, while the clustering isolating MSE was based on difference ERPs resulting from the subtraction of D_AV_ and D_V_. Thus, 4 group-averaged ERPs were submitted to both clustering analyses, one for each of the context-related conditions. Next, the data were fitted onto the canonical N2pc time-window (180–300ms) as well as other, earlier time-periods, notably, also ones including time-periods highlighted by the GFP results as representing significant condition differences. The resulting map presence (in ms) over the given time-windows were submitted to 4-way rmANOVAs with factors: Multisensory Relationship (Arbitrary vs. Congruent), Distractor Onset (Unpredictable vs. Predictable), and Map (different numbers of maps, depending on the topographic clustering analyses and time-windows within each clustering analyses), followed by post-hoc *t*-tests. Maps with durations <15 contiguous timeframes were not included in the analyses. Unless otherwise stated in the Results, map durations were statistically different from 0ms (as confirmed by post-hoc one-sample t-tests), meaning that they were reliably present across the time-windows of interest. Holm-Bonferroni corrections (Holm, 1979) were used to correct for multiple comparisons between map durations. Comparisons passed the correction unless otherwise stated.

## Results

### Behavioural analyses

#### Interaction of TAC and MSE with contextual factors

To shed light on attentional control in naturalistic settings, we first tested whether top-down visual control indexed by TAC interacted with contextual factors in behavioural measures. First, our 2 × 2 × 2 × 2 rmANOVA confirmed the presence of TAC, via a main effect of Distractor Colour, *F*_(1, 38)_ = 340.4, *p* < 0.001, η_p_^2^ = 0.9, with TCC distractors (42ms), but not NCC distractors (−1ms), eliciting reliable behavioural capture effects. Of central interest here, the strength of TAC was dependent on whether the multisensory relationship within the distractor involved mere simultaneity or semantic congruence. This was demonstrated by a 2-way Distractor Colour × Multisensory Relationship interaction, *F*_(1, 38)_ = 4.5, *p* = 0.041, η_p_^2^ = 0.1 (Figure 2). This effect was driven by behavioural capture effects elicited by TCC distractors being reliably larger for the Arbitrary (45ms) than for the Congruent (40ms) condition, *t*_(38)_ = 1.9, *p* = 0.027. NCC distractors showed no evidence of Multisensory Relationship modulation (Arbitrary vs. Congruent, *t*_(38)_ = 1, *p* = 0.43). Contrastingly, TAC showed no evidence of modulation by predictability of the distractor onset (no 2-way Distractor Colour × Distractor Onset interaction, *F*_(1, 38)_ = 2, *p* = 0.16). Thus, visual feature-based attentional control interacted with the contextual factor of distractor semantic congruence, but not distractor temporal predictability.

**Figure 2.**
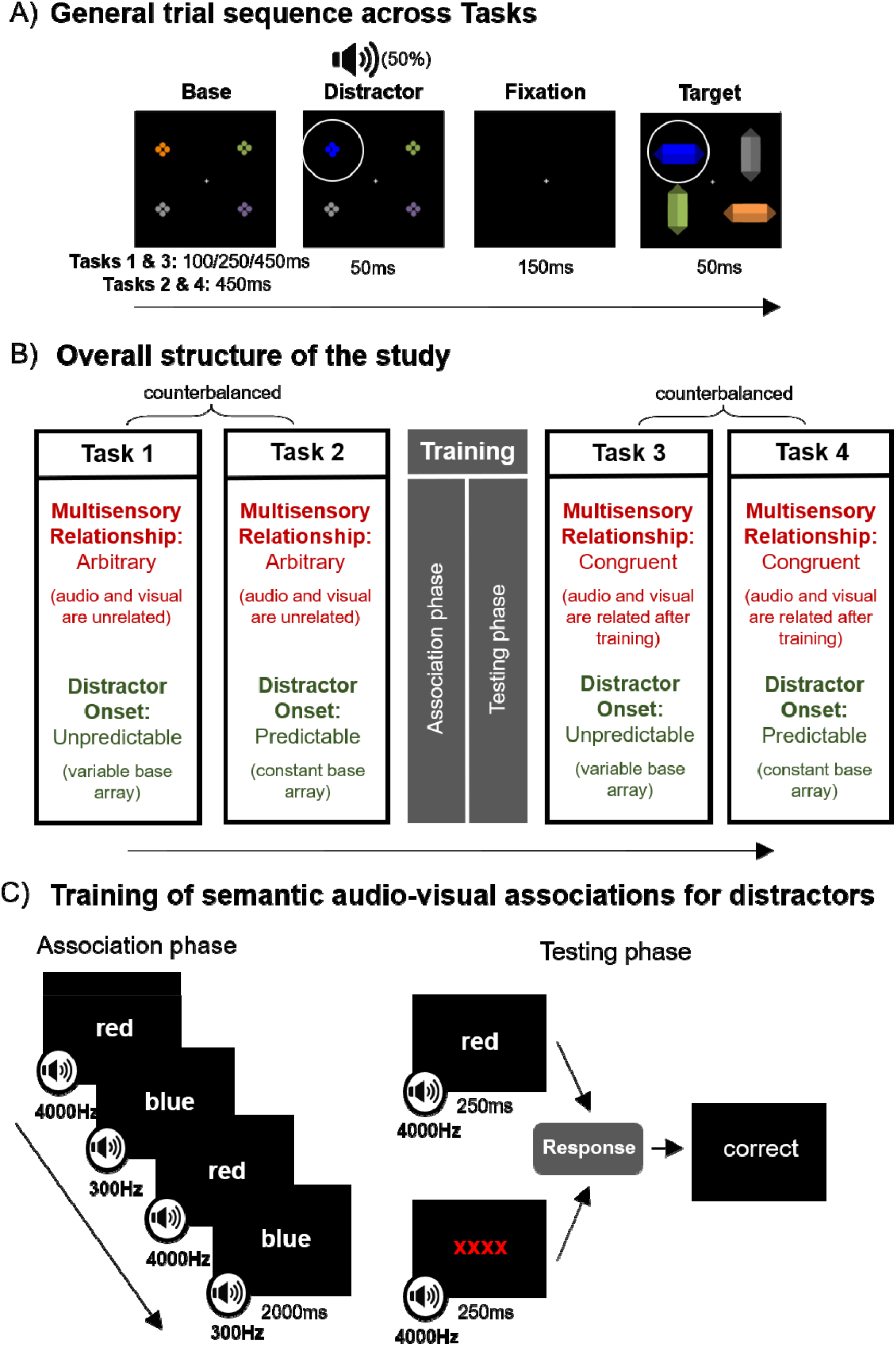
The violin plots show the attentional capture effects (spatial cueing in milliseconds) for TCC and NCC distractors, and the distributions of single-participant scores according to whether Multisensory Relationship within these distractors was Arbitrary (light green) or Congruent (dark green). The dark grey boxes within each violin plot show the interquartile range from the 1^st^ to the 3^rd^ quartile, and white dots in the middle of these boxes represent the median. Larger values indicate *positive* behavioural capture effects (RTs faster on trials where distractor and target appeared in same vs. different location), while below-zero values – *inverted* capture effects (RTs slower on trials where distractor and target appeared in same vs. different location). Larger behavioural capture elicited by target-colour distractors (TCC) was found for arbitrary than semantically congruent distractors. Expectedly, regardless of Multisensory Relationship, attentional capture was larger for target-colour (TCC) distractors than for non-target colour distractors (NCC).

Next, we investigated potential interactions of multisensory enhancements with contextual factors. Expectedly, there was behavioural MSE (a significant main effect of Distractor Modality, *F*_(1, 38)_=13.5, *p*=0.001, η_p_^2^=0.3), where visually-elicited behavioural capture effects (18ms) were enhanced on AV trials (23ms). Unlike TAC, this MSE effect showed no evidence of interaction with either of the two contextual factors (Distractor Modality x Multisensory Relationship interaction, *F*<1; Distractor Modality x Distractor Onset interaction: *n.s*. trend, *F*_(1, 38)_=3.6, *p*=0.07, η_p_^2^= 0.1). Thus, behaviourally, Multisensory enhancement of attentional capture was not modulated by distractors’ semantic relationship nor its temporal predictability. We have also observed other, unexpected effects, but as these were outside of the focus of the current paper, which aims to elucidate the interactions between visual (goal-based) and multisensory (salience-driven) attentional control and contextual mechanisms, we describe them only in SOMs.

### ERP analyses

#### Lateralised (N2pc-like) brain mechanisms

We next investigated the type of brain mechanisms that underlie interactions between more traditional attentional control (TAC, MSE) and contextual control over attentional selection. Our analyses on the lateralised responses, spanning both a canonical and EN framework, revealed little evidence for a role of spatially-selective mechanisms in supporting the above interactions. Both canonical N2pc and electrical neuroimaging analyses confirmed the presence of TAC (see Fig. 3 for N2pc waveforms across the four distractor types). However, TAC did not interact with either of the two contextual factors. Lateralised ERPs also showed no evidence for sensitivity to MSE nor for interactions between MSE and any contextual factors. Not even the main effects of Multisensory Relationship and Distractor Onset^6^ were present in lateralised responses (See SOMs for full description of the results of lateralised ERP analyses).

**Figure 3.**
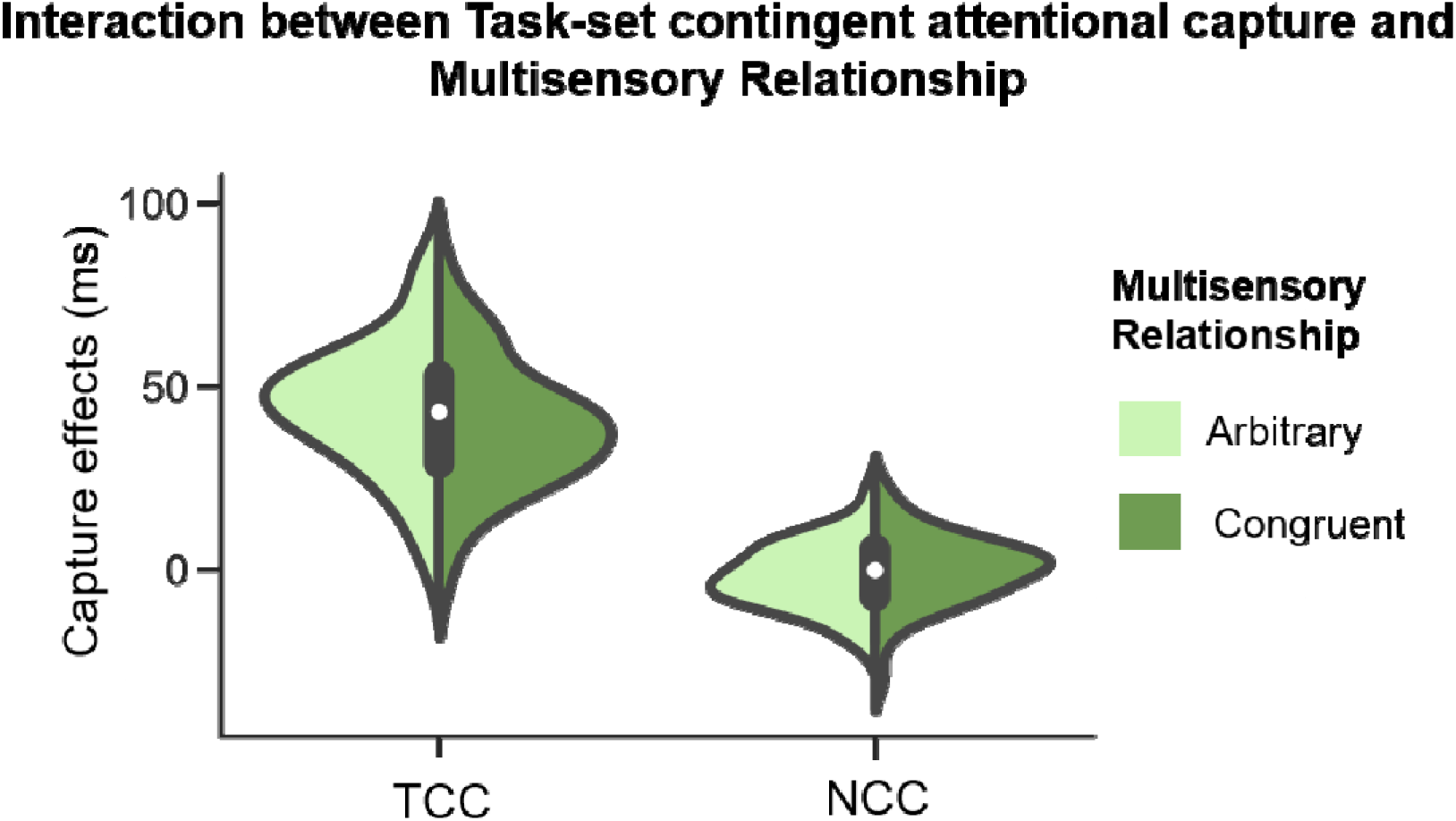
Overall contra- and ipsilateral ERP waveforms representing a mean amplitude over electrode clusters (plotted on the head model at the bottom of the figure in blue and black), separately for each of the four experimental conditions (Distractor Colour x Distractor Modality), averaged across all four Tasks. The N2pc time-window of 180–300ms following distractor onset is highlighted in grey, and significant contra-ipsi differences are marked with an asterisk (*p* < 0.05). As expected, only the TCC distractors elicited statistically significant contra-ipsi differences.

#### Nonlateralised brain mechanisms

A major part of our analyses focused on understanding the role of nonlateralised ERP mechanisms in the interactions between visual goals (TAC), multisensory salience (MSE) and contextual control. To remind the reader, to prevent nonlateralised ERPs from being confounded by the presence of sound on AV trials, we based our analyses here on the difference ERPs indexing visual attentional control under sound absence vs. presence. That is, we calculated ERPs of the difference between TCCV and NCCV conditions, and between TCCAV and NCCAV conditions (D_V_ and D_AV_ levels, respectively, of the Target Difference factor). We focus the description of these results on the effects of interest (see SOMs for full description of results).

The 2 × 2 × 2 (Multisensory Relationship × Distractor Onset × Target Difference) rmANOVA on electrode-wise voltage analyses revealed a main effect of Target Difference at 53–99ms and 141–179ms, thus both at early, perception-related, and later, attentional selection-related latencies (reflected by the N2pc). Across both time-windows, amplitudes were larger for D_AV_ (TCCAV – NCCAV difference) than for D_V_ (TCCV – NCCV difference). This effect was further modulated, evidenced by a 2-way Target Difference × Multisensory Relationship interaction, at the following time-windows: 65–103ms, 143–171ms, and 194–221ms (all *p*’s < 0.05). The interaction was driven by Congruent distractors showing larger amplitudes for D_AV_ than D_V_ within all 3 time-windows (65–97ms, 143–171ms, and 194–221ms; all *p*’s < 0.05). No similar differences were found for Arbitrary distractors, and there were no other interactions that passed the temporal and spatial criteria for multiple comparisons of >15 contiguous timeframes and >10% of the 129-channel electrode montage.

#### Interaction of TAC with contextual factors

We next used electrical neuroimaging analyses to investigate the contribution of the strength- and topography-based nonlateralised mechanisms to the interactions between TAC and contextual factors.

##### Strength-based brain mechanisms

A 2 × 2 × 2 Target Difference × Multisensory Relationship × Distractor Onset rmANOVA on the GFP mirrored the results of the electrode-wise analysis on ERP voltages by showing a main effect of Target Difference spanning a large part of the first 300ms post-distractor both before and in N2pc-like time-windows (19–213ms, 221–255ms, and 275–290ms). Like in the voltage waveform analysis, the GFP was larger for D_AV_ than D_V_ (all *p*’s < 0.05). In GFP, Target Difference interacted both with Multisensory Relationship (23–255ms) and separately with Distractor Onset (88–127ms; see SOMs for full description). Notably, there was a 3-way Target Difference × Multisensory Relationship × Distractor Onset interaction, spanning 102–124ms and 234–249ms. We followed up this interaction with a series of post-hoc tests to gauge the modulations of TAC (and MSE, see below) by the two contextual factors.

In GFP, Multisensory Relationship and Distractor Onset interacted independently of Target Difference in the second time-window, which results we describe in SOMs. To gauge differences in the strength of TAC in GFP across the 4 contexts (i.e., Arbitrary Unpredictable, Arbitrary Predictable, Congruent Unpredictable, and Congruent Predictable), we focused the comparisons on only visually-elicited target differences (to minimise any potential confounding influences from sound processing) across the respective levels of the 2 contextual factors. The weakest GFPs were observed for Arbitrary Predictable distractors (Figure 4A). They were weaker than GFPs elicited for Arbitrary Unpredictable distractors (102–124ms and 234–249ms), and Predictable Congruent distractors (only in the later window, 234–249ms).

**Figure 4.**
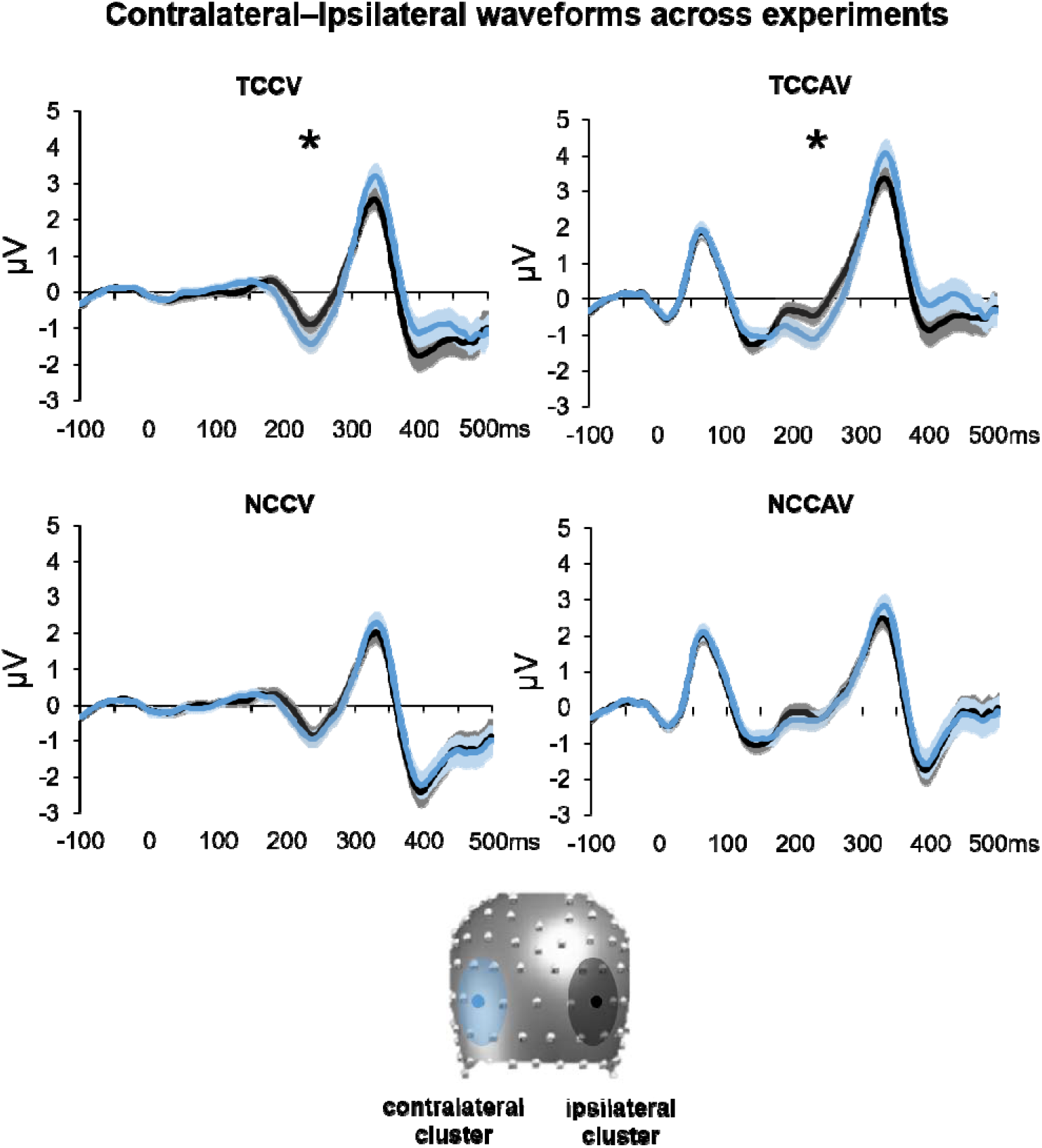
Nonlateralised GFP and topography results for the visual only difference ERPs (DV condition of Target Difference), as a proxy for TAC. **A)** Mean GFP over the post-distractor and pre-target time-period across the 4 experimental tasks (as a function of the levels of Multisensory Relationship and Distractor Onset that they represent), as denoted by the colours on the legend. The time-windows of interest (102–124ms and 234–249ms) are highlighted by grey areas. **B)** Template maps over the post-distractor time-period as revealed by the topographic clustering (Maps A1 to A5) are shown in top panels. In lower panels are the results of the fitting procedure over the 29–126ms time-window. The results displayed here are the follow-up tests of the 3-way Map x Multisensory Relationship x Distractor Onset interaction as a function of Multisensory Relationship (leftward panel) and of Distractor Onset (rightward panel). Bars are coloured according to the template maps that they represent. Conditions are represented by full colour or patterns per the legend. Error bars represent standard errors of the mean.

##### Topography-based brain mechanisms

We focused the topographic clustering of the TAC-related topographic activity on the whole 0–300ms post-distractor time-window (before the target onset), which revealed 10 clusters that explained 82% of the global explained variance within the visual-only ERPs. This time-window of 29–126ms post-distractor was selected on based on the GFP peaks, which are known to correlate with topographic stability (Lehmann 1987; Brunet et al. 2011), and in some conditions, based on the fact that specific template was dominated responses in group-averaged data from given conditions, e.g., Arbitrary Unpredictable and Congruent Unpredictable conditions, but not for other conditions. This was confirmed by our statistical analyses, with a 2 × 2 × 5 rmANOVA over the 29–126ms post-distractor time-window, which revealed a 3-way Multisensory Relationship × Distractor Onset × Map interaction, *F*_(3.2,122)_ = 5.3, *p* = 0.002, η_p_^2^ = 0.1.

Follow-up tests in the 29–126ms time-window focused on maps differentiating between the 4 contexts as a function of the two contextual factors (results of follow-up analyses as a function of Multisensory Relationship and Distractor Onset are visible in Figure 4B in leftward panel and rightward panel, respectively). These results confirmed that context altered the processing of distractors from early on. The results also confirmed the clustering that the context did so by engaging different networks for most of the different combinations of Multisensory Relationship and Distractor Onset: Arbitrary Unpredictable - Map A2, Congruent Unpredictable - Map A5, as well as for Arbitrary Predictable - Map A1 (no map predominantly involved in the responses for Congruent Predictable).

*Arbitrary Predictable distractors*, which elicited the weakest GFP, recruited predominantly Map A1 (37ms) during processing. This map was more involved in the processing of those distractors vs. Congruent Predictable distractors (21ms), *t*_(38)_ = 2.7, *p* = 0.013 (Fig. 4B bottom panel).

*Arbitrary Unpredictable* distractors largely recruited Map A2 (35ms) during processing. This map was more involved in the processing of these distractors vs. Arbitrary Predictable distractors (18ms), *t*_(38)_ = 2.64, *p* = 0.012 (Fig. 4B top leftward panel), as well as Congruent Unpredictable distractors (14ms), *t*_(38)_ = 3.61, *p* < 0.001 (Fig. 4B top rightward panel).

*Congruent Unpredictable* distractors principally recruited Map A5 (34ms) during processing, which was more involved in the processing of these distractors vs. Congruent Predictable distractors (19ms) distractors, *t*_(38)_ = 2.7, *p* = 0.039 (Fig. 4B middle leftward panel), as well as Arbitrary Unpredictable (12ms) distractors, *t*_(38)_ = 3.7, *p* <0.001 (Fig. 4B middle rightward panel).

*Congruent Predictable* distractors recruited different template maps during processing, where Map A2 was more involved in responses to those distractors (25ms) vs. Congruent Unpredictable distractors (14ms), *t*_(38)_ = 2.17, *p* = 0.037, but not other distractors, *p*’s>0.2 (Fig. 4B top leftward panel).

#### Interaction of MSE with contextual factors

We next analysed the strength- and topography-based nonlateralised mechanisms contributing to the interactions between MSE and contextual factors.

##### Strength-based brain mechanisms

To gauge the AV-induced enhancements between D_AV_ and D_V_ across the 4 contexts, we explored the abovementioned 2 × 2 × 2 GFP interaction using a series of simple follow-up post-hoc tests. We first tested if response strength between D_AV_ and D_V_ was reliably different within each of the 4 contextual conditions. AV-induced ERP responses were enhanced (i.e., larger GFP for D_AV_ than D_V_ distractors) for both Predictable and Unpredictable Congruent distractors, across both earlier and later time-windows. Likewise, AV enhancements were also found for Arbitrary Predictable distractors, but only in the earlier (102–124ms) time-window. Unpredictable distractors showed similar GFP across D_AV_ and D_V_ trials. Next, we compared the AV-induced MS enhancements across the 4 contexts, by creating (D_AV_ minus D_V_) difference ERPs or each context. AV-induced enhancements were weaker for Predictable Arbitrary distractors than Predictable Congruent distractors (102–124ms and 234–249ms; Figure 5A).

**Figure 5.**
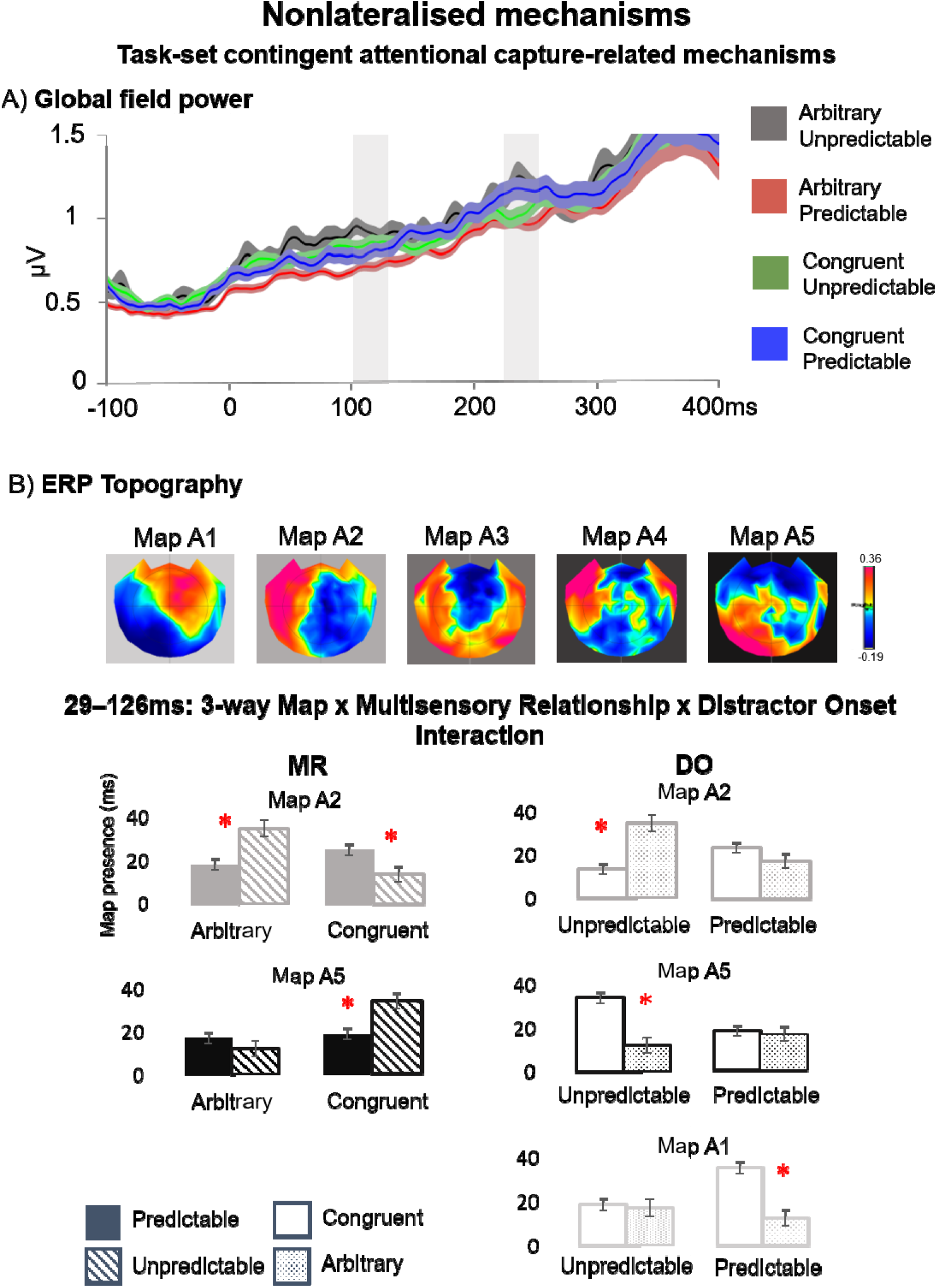
Nonlateralised GFP and topography results for the difference ERPs between the DAV and DV conditions of Target Difference, as a proxy for MSE. **A)** Mean GFP over the postdistractor and pre-target time-period across the 4 experimental tasks (as a function of the levels of Multisensory Relationship and Distractor Onset that they represent), as denoted by the colours on the legend. The time-windows of interest (102–124ms and 234–249ms) are highlighted by grey bars. **B)** Template maps over the post-distractor time-period as revealed by the topographic clustering (Maps A1 to A7) are shown on top. Below are the results of the fitting procedure over the three time-windows: 35–110, 110–190, and 190–300 time-window. Here we display the follow-ups of the interactions observed in each time-window: in 35–110 and 190–300 time-windows, the 2-way Map x Multisensory Relationship interaction (leftward and rightward panels, respectively), and in the 110–190 time-window, follow-ups of the 3-way Map x Multisensory Relationship x Distractor Onset interaction as a function of Multisensory Relationship and of Distractor Onset (middle panel). Bars are coloured according to the template maps that they represent. Conditions are represented by full colour or patterns per the legend. Error bars represent standard errors of the mean.

Topography-based brain mechanisms. We then used the difference (D_AV_ minus D_V_) difference ERPs (as in the second part of the GFP analyses) to focus the topographic clustering selectively on the MSE-related topographic activity. This clustering, carried out on the 0–300ms post-distractor and pre-target time-window, revealed 7 clusters that explained 78% of the global explained variance within the AV-V target difference ERPs.

In this topographic clustering there were multiple GPF peaks, with elongated near-synchronous periods of time where different maps were suggested to be present across the four distractor conditions in the group-averaged data. One of those maps (Map B3) was first present in the two congruent distractor conditions, to then become absent and reappear again. In the view of this patterning, we decided to fit the group-average data from these three subsequent time-windows to single-subject data: 35–110ms, 110–190ms, and 190–300ms. To foreshadow the results, in the first and third time-windows the MSE-related template maps were modulated only by Multisensory Relationship, while in the middle time-window – by both Multisensory Relationship and Distractor Onset.

In the first, 35–110ms time-window, the modulation of map presence by Multisensory Relationship was evidenced by a 2-way Map × Multisensory Relationship interaction, *F*_(2.1,77.9)_ = 9.2, *p* < 0.001, η_p_^2^ = 0.2. This effect was driven by one map (map B3) that, in this time-window, predominated responses to Congruent (42ms) vs. Arbitrary (25ms) distractors, *t*_(38)_= 4.3, *p* = 0.02, whereas another map (map B5) dominated responses to Arbitrary (33ms) vs. Congruent (18ms) distractors, *t*_(38)_ = 4, *p* = 0.01 (Figure 5B top and upper leftward panels, respectively).

In the second, 110–190ms time-window, map presence was modulated by both contextual factors, with a 3-way Map × Multisensory Relationship × Distractor Onset interaction, *F*_(2.6,99.9)_ = 3.7, *p* = 0.02, η_p_^2^ = 0.1 (just as it did for TAC). We focused follow-up tests in that time-window again on maps differentiating between the 4 conditions, as we did for the 3-way interaction for TAC (results of follow-ups as a function of Multisensory Relationship and Distractor Onset are visible in Figure 5B, middle upper and lower panels, respectively). Context processes again interacted to modulate the processing of distractors, although now they did so after the first 100ms. They did so again by engaging different networks for different combinations of Multisensory Relationship and Distractor Onset: Arbitrary Predictable distractors - Map B1, Arbitrary Unpredictable distractors - Map B5, Congruent Unpredictable distractors - Map B6, and now also Congruent Predictable distractors - Map B3.

*Arbitrary Predictable* distractors, which again elicited the weakest GFP, during processing mainly recruited map B1 (35ms). This map dominated responses to these distractors vs. Arbitrary Unpredictable distractors (18ms, *t*_(38)_ = 2.8, *p* = 0.01; Figure 5B upper panel), as well as Congruent Predictable distractors (17ms, *t*_(38)_ = 2.8, *p* = 0.006; Figure 5B lower panel).

*Arbitrary Unpredictable* distractors largely recruited during processing one map, Map B5 (33ms). Map B5 was more involved in responses to these distractors vs. Arbitrary Predictable distractors (17ms, *t*_(38)_ = 2.6, *p* = 0.042; Figure 5B upper panel), as well as vs. Congruent Unpredictable distractors (13ms, *t*_(38)_ = 3.4, *p* = 0.002; Figure 5B bottom panel).

*Congruent Unpredictable* distractors principally recruited during processing Map B6 (37ms). Map B6 was more involved in responses to these distractors vs. Congruent Predictable distractors (21ms, *t*_(38)_ = 2.5, *p* = 0.02), and vs. Arbitrary Unpredictable distractors (24ms, *t*_(38)_= 2.3, *p* = 0.044).

*Congruent Predictable* distractors mostly recruited during processing Map B3 (25ms). Map B3 was more involved in responses to these distractors vs. Predictable Arbitrary distractors (8ms, *t*_(38)_ = 2.2, *p* = 0.005), and, at statical-significance threshold level, vs. Congruent Unpredictable distractors (12ms, *t*_(38)_ = 2.2, *p* = 0.0502).

In the third, 190–300ms time-window, the 2-way Map × Multisensory Relationship interaction was reliable at *F*_(3.2,121.6)_ = 3.7, *p* = 0.01, η_p_^2^= 0.1. Notably, the same map as before (map B3) was more involved, at a non-statistical trend level, in the responses to Congruent (50ms) vs. Arbitrary distractors (33ms), *t*_(38)_ = 3.6, *p* = 0.08, and another map (map Bl) predominated responses to Arbitrary (25ms) vs. Congruent (14ms) distractors, *t*_(38)_ = 2.3, *p* = 0.02 (Figure 5B rightward panel).

## Discussion

Attentional control is necessary to cope with the multitude of stimulation in everyday situations. However, in such situations, the observer’s goals and stimuli’s salience routinely interact with contextual processes, yet such multi-pronged interactions between control processes have never been studied. Below, we discuss our findings on how visual and multisensory attentional control interact with distractor temporal predictability and semantic relationship. We then discuss the spatiotemporal dynamics in nonlateralised brain mechanisms underlying these interactions. Finally, we discuss how our results enrich the understanding of attentional control in real-world settings.

### Interaction of task-set contingent attentional capture with contextual control

Visual control interacted most robustly with stimuli’s semantic relationship. Behaviourally, *target-matching* visual distractors captured attention more strongly when they were arbitrarily connected than semantically congruent. This was accompanied by a cascade of modulations of nonlateralised brain responses, spanning both the attentional selection, N2pc-like stage and much earlier, perceptual stages. Arbitrary distractors, but only predictable ones, first recruited one particular brain network (Map A1), to a larger extent than predictable semantically congruent distractors, and did so early on (29–126ms post-distractor). Arbitrary predictable distractors elicited also suppressed responses, in the later part of this early time-window (102–124ms; where they elicited the weakest responses). In the later, N2pc-like (234–249ms) time-window, responses to arbitrary predictable distractors were again weaker, now compared to semantically congruent predictable distractors.

This cascade of network- and strength-based modulations of nonlateralised brain responses might epitomise a potential brain mechanism for interactions between visual topdown control and multiple sources of contextual control, as they are consistent with existing literature. The discovered early (~30-100ms) topographic modulations for predictable target-matching (compared to unpredictable) distractors is consistent with predictions attenuating the earliest visual perceptual stages (C1 component, ~50–100ms post-stimulus; Dassanayake et al. 2016). The subsequent, mid-latency response suppressions (102–124ms, where we found also topographic modulations) for predictable distractors are in line with N1 attenuations for self-generated sounds (Baess et al. 2011; Klaffehn et al. 2019), and the latencies where the brain might promote the processing of unexpected events (Press et al. 2020). Notably, these latencies are also in line with the onset (~115ms post-stimulus) of the goal-based suppression of salient visual distractors (here: presented simultaneously with targets), i.e., distractor positivity (Pd; Sawaki & Luck 2010). Finally, the response suppressions we found at later, N2pc-like, attentional selection stages (234–249ms), are also consistent with some extant (albeit scarce) literature. Van Moorselaar and Slagter (2019) showed that when such salient visual distractors appear in predictable locations, they elicit the N2pc but no longer a (subsequent, post-target) Pd, suggesting that once the brain learns the distractor’s location, it can suppress it without the need for active inhibition. More recently, van Moorselaar et al. (2020b) showed that the representation of the predictable distractor feature could be decoded already from pre-stimulus activity. While our paradigm was not optimised for revealing such effects, pre-stimulus mechanisms could indeed explain our early-onset (~30ms) context-elicited neural effects. The robust response suppressions for predictable stimuli are also consistent with recent proposals for interactions between predictions and auditory attention. Schröger et al. (2015) suggested that greater attention is deployed to more “salient” stimuli, i.e., those for which a prediction is missing, so that the predictive model can be reconfigured to encompass such predictions in the future. This reconfiguration, in turn, requires top-down goal-based attentional control. Our results extend this model to the visual domain. Our findings involving the response modulation cascade and behavioural benefits may also support the Schröger et al.’s tenet that different, but connected, predictive models exist at different levels of the cortical hierarchy.

These existing findings jointly strengthen our interpretations that goal-based topdown control utilises contextual information to alter visual processing from very early on in life. Our findings also extend the extant ideas in several ways. First, they show that in context-rich settings (i.e., involving multiple sources of contextual control), goal-based control will use both stimulus-related predictions and stimulus meaning to facilitate task-relevant processing. Second, context information modulates not only early, pre-stimulus and late, attentional stages, but also early *stimulus-elicited* responses. Third, our findings also suggest candidate mechanisms for supporting interactions between goal-based control and multiple sources of contextual information. Namely, context will modulate the early stimulus processing by recruiting distinct brain networks for stimuli representing different contexts, e.g., the brain networks recruited by predictable distractors differed for arbitrarily linked and semantically congruent stimuli (Map A1 and A2, respectively). Also, the distinct network recruitment might lead to the suppressed (potentially more efficient; c.f. repetition suppression, Grill-Spector et al. 2006) brain responses. These early response attenuations will extend also to later stages, associated with attentional selection. Thus, it is the early differential brain network recruitment that might trigger a cascade of spatiotemporal brain dynamics leading effectively to the stronger behavioural capture, here for predictable (arbitrary) distractors. However, for distractors, these behavioural benefits may be most robust for arbitrary target-matching stimuli (as opposed to semantically congruent), with prediction-based effects are less apparent.

### Interaction of multisensory enhancement of attentional capture with contextual control

Across brain responses, multisensory-induced processes interacted with both contextual processes. To measure effects related to multisensory-elicited modulations and to its interactions with contextual information, we analysed AV–V differences within the Target Difference ERPs.

The interactions between multisensory modulations and context processes were also instantiated via an early-onset cascade of strength- and topographic (network-based) nonlateralised brain mechanisms. This cascade again started early (now 35–110ms post-distractor). A separate topographic clustering analysis revealed that in the multisensory-modulated responses the brain first distinguished only between semantically congruent and arbitrarily linked distractors. These distractors recruited predominantly different brain networks (Map B3 and B5, respectively). Around the end of these topographic, network-based modulations, at 102–124ms, multisensory-elicited brain responses were also modulated in their strength. Arbitrary predictable distractors again triggered weaker responses, now compared to semantically congruent predictable distractors. Multisensory-elicited responses predominantly recruited distinct brain networks for the four distractor types from 110ms until 190ms post-distractor, thus spanning stages linked to perception and attentional selection. Here, maps B3 and B5 were now recruited for responses to semantically congruent predictable and arbitrary unpredictable distractors, respectively. Meanwhile, maps B1 and B6 were recruited for arbitrary predictable and semantically congruent unpredictable distractors, respectively. In the subsequent time-window (190–300ms) that mirrors the time-window used in the canonical N2pc analyses, multisensory-related responses again recruited different brain networks. There, Map B3 (previously: Congruent Predictable distractors) again was predominantly recruited by semantically congruent over arbitrary distractors, and now Map B1 (previously: Arbitrary Predictable distractors) - for arbitrary distractors over congruent ones. In the middle of this time-window (234–249ms), responses differed in their strength, with predictable arbitrary distractors eliciting weaker responses compared to semantically congruent predictable distractors.

To summarise, distractors’ semantic relationship played a dominant (but not absolute) role in interactions between multisensory-elicited and contextual processes. The AV-V difference ERPs were modulated exclusively by multisensory relationships both in the earliest, perceptual (35–110ms) time-window and latest, N2pc-like (190–300ms) time-window linked to attentional selection. At both stages, distinct brain networks were recruited predominantly by semantically congruent and arbitrary distractors. These results suggest that from early perceptual stages the brain “relays” the processing of (multisensory) stimuli as a function of them containing meaning (vs. lack thereof) for the observer up to stages of attentional selection. Notably, the same brain network (Map B3) supported multisensory processing of semantically congruent distractors across both time-windows, while different networks were recruited by arbitrarily linked distractors.

Thus, a single network might be recruited for processing meaningful multisensory stimuli. In light of our behavioural results, this brain network could be involved in suppressing behavioural attentional capture for semantically congruent (over arbitrarily linked) distractors by top-down goal-driven attentional control. This idea is supported by the interactions between distractors’ multisensory-driven modulations, their multisensory relationship, and their temporal predictability in the second, 110–190ms time-window. Therein, the same “semantic” Map B3 was still present, albeit now recruited for responses to semantically congruent (over arbitrary) *predictable* distractors. Based on existing evidence that predictions are used in service of goal-based behaviour (Schröger et al. 2015; van Moorselaar et al. 2020a; Matusz et al. 2016), one could argue that the brain network reflected by Map B3 might play a role in integrating contextual information across both predictions and meaning (though mostly meaning, as it remained recruited by semantically congruent distractors throughout the distractor-elicited response). The activity of this network might have contributed to the overall stronger brain responses (indicated by GFP results) to semantically congruent multisensory stimuli, which in turn contributed to the null behavioural multisensory enhancements of behavioural indices of attentional capture. While these are the first results of this kind, they open an exciting possibility that surface-level EEG/ERP studies can reveal the network- and strength-related brain mechanisms (potentially a single network for “gain control” up-modulation) by which goal-based processes control (i.e., suppress) multisensorily-driven enhancements of attentional capture.

### Towards understanding how we pay attention in naturalistic settings

It is now relatively well-established that the brain facilitates goal-directed processing (from perception to attentional selection) via processes based on observer’s goals (e.g. Folk et al. 1992; Desimone & Duncan 1995), predictions about the outside world (Summerfield & Egner 2009; Schröger et al. 2015; Press et al. 2020), and long-term memory contents (Summerfield et al. 2006; Peelen & Kastner 2014). Also, multisensory processes are increasingly recognised as an important source of bottom-up, attentional control (e.g. Spence & Santangelo 2007; Matusz & Eimer 2011; Matusz et al. 2019a; Fleming et al. 2020). By studying these processes largely in isolation, researchers clarified how they support goal-directed behaviour. However, in the real world, observer’s goals interact with multisensory processes and multiple types of contextual information. Our study sheds first light on this “naturalistic attentional control”.

Understanding of attentional control in the real world has been advanced by research on feature-related mechanisms (Theeuwes 1991; Folk et al. 1992; Desimone & Duncan 1995; Luck et al. 2020), which support attentional control where target location information is missing. Here, we aimed to increase the ecological validity of this research by investigating how visual feature-based attention (as indexed by TAC) transpires in context-rich, multisensory settings (see SOMs for a discussion of our replication of TAC). Our findings of reduced capture for semantically congruent than artificially linked target-colour matching distractors is novel and important, as they suggest stimuli’s meaning is also utilised to suppress attention (to distractors). Until now, known benefits of meaning were limited to target selection (Thorpe et al. 1996; Iordanescu et al. 2008; Matusz et al. 2019a). Folk et al. (1992) famously demonstrated that attentional capture by distractors is sensitive to the observer’s goals; we reveal that distractor’s meaning may serve as a second source of goal-based attentional control. This provides a richer explanation for how we stay focused on task in everyday situations, despite many objects matching attributes of our current behavioural goals.

To summarise, in the real world, attention should be captured more strongly by stimuli that are unpredictable (Schröger et al. 2015), but also by those unknown or without a clear meaning. On the other hand, stimuli with high strong spatial and/or temporal alignment across senses (and so stronger bottom-up salience) may be more resistant to such goal-based attentional control (suppression), as we have shown here (multisensory enhancement of attentional capture; see also Santangelo & Spence 2007; Matusz & Eimer 2011; van der Burg et al. 2011; Turoman et al. 2021a; Fleming et al. 2020). As multisensory distractors captured attention more strongly even in current, context-rich settings, this confirms the importance of multisensory salience as a source of *potential* bottom-up attentional control in naturalistic environments (see SOMs for a short discussion of this replication).

The investigation of brain mechanisms underlying known EEG/ERP correlates (N2pc, for TAC) via advanced multivariate analyses has enabled us to provide a comprehensive, novel account of attentional control in a multi-sensory, context-rich setting. Our results jointly support the primacy of goal-based control in naturalistic settings. Multisensory semantic congruence reduced behavioural attentional capture by target-matching colour distractors compared to arbitrarily linked distractors. Context modulated nonlateralised brain responses to target-related (TAC) distractors via a cascade of strength- and topographic mechanisms from early (~30ms post-distractor) to later, attentional selection stages. While these results are first of this kind and need replication, they suggest that context-based goal-directed modulations of distractor processing “snowball” from early stages (potentially involving pre-stimulus processes, e.g. van Moorselaar & Slagter, 2020) to control behavioural attentional selection. Responses to predictable arbitrary (target-matching) distractors revealed by our electrical neuroimaging analyses might have driven the larger behavioural capture for arbitrary than semantically congruent distractors. The former engaged distinct brain networks and triggered the weakest and potentially most efficient (Grill-Spector et al. 2006) responses. One reason for the absence of such effects in behavioural measures is the small magnitude of behavioural effects: while the TAC effect is ~50ms, both MSE effect and semantically-driven suppression were small, at around ~5ms. This may also be the reason why context-driven effects were absent in behavioural measures of multisensory enhancement of attentional capture, despite involving a complex, early-onsetting cascade of strength- and topographic modulations.

Our results point to a potential brain mechanism by which semantic relationships influence goal-directed behaviour towards task-irrelevant information. Namely, our electrical neuroimaging analyses of surface-level EEG identified a brain network that is recruited by semantically congruent stimuli at early, perceptual stages, and that remains active at N2pc-like, attentional selection stages. While remaining cautious when interpreting our results, this network might have contributed to the consistently enhanced AV-induced responses for semantically congruent multisensory distractors. These enhanced brain responses together with the concomitant *suppressed behavioural attention* effects are consistent with a “gain control” mechanism, in the context of distractor processing (e.g. Sawaki & Luck 2010; Luck et al. 2020). Our results reveal that such “gain control”, at least in some cases, operates by relaying processing of certain stimuli to distinct brain networks. We have purported the existence of such a “gain control” mechanism in a different study on (top-down) multisensory attention (e.g. Matusz et al. 2019c). While these are merely speculations that would require source estimations to be supported, the enhanced responses to meaningful distractors may thus reflect enhanced goal-based control over those stimuli. Such a process could potentially recruit a network involving the anterior hippocampus and putamen, which help maintain active representations of task-relevant information while updating the representation of to-be-suppressed information (McNab & Klingberg 2008; Sadeh et al. 2010; Jiang et al. 2015). Our electrical neuroimaging analyses of the surface-level N2pc data (see also Matusz et al. 2019c; Turoman et al. 2021a) might have potentially revealed when and how such memory-related brain networks modulate attentional control over task-irrelevant stimuli.

### N2pc as an index of attentional control

We have previously discussed the limitations of canonical N2pc analyses in capturing neurocognitive mechanisms by which visual top-down goals and multisensory bottom-up salience simultaneously control attention selection (Matusz et al. 2019b). The mean N2pc amplitude modulations are commonly interpreted as “gain control”, but they can be driven by both strength- (i.e., “gain”) and topographic (network-based) mechanisms. Canonical N2pc analyses cannot distinguish between those two brain mechanisms. Contrastingly, Matusz et al. (2019b) have shown evidence for both brain mechanisms underlying N2pc-like responses. These and other results of ours (Turoman et al. 2021a, 2021b) provided evidence from surface-level data for different brain sources contributing to the N2pc’s, a finding that has been previously shown only in source-level data (Hopf et al. 2000). These findings point to a certain limitation of the N2pc (canonically analysed), which is an EEG *correlate* of attentional selection, but where other analytical approaches are necessary to reveal brain mechanisms of attentional selection.

Here, we have shown that the lateralised, spatially-selective brain mechanisms, approximated by the N2pc and revealed by electrical neuroimaging analyses are limited in how they contribute to attentional control in some settings. Rich, multisensory, and context-laden influences over goal-based top-down attention are, in our current paradigm, not captured by such lateralised mechanisms. In contrast, nonlateralised (or at least *relatively less* lateralised, see Figures 4 and 5) brain networks seem to support such interactions for visual and multisensory distractors - from early on, leading to attentional selection. We nevertheless want to reiterate that paradigms that can gauge N2pc offer an important starting point for studying attentional control in less traditional multisensory and/or context-rich settings. There, multivariate analyses, and an electrical neuroimaging framework in particular, might be useful in readily revealing new mechanistic insights into attentional control.

### Broader implications

Our findings are important to consider when aiming to study attentional control, and information processing more generally, in naturalistic settings (e.g., while viewing movies, listening to audiostories) and veridical real-world environments (e.g. the classroom or the museum). Additionally, conceptualisations of ecological validity (Peelen et al. 2014; Shamay-Tsoory & Mendelsohn 2019; Vanderwal et al. 2019; Eickhoff et al. 2020; Cantlon 2020) should go beyond traditionally invoked components (e.g., observer’s goals, context, socialness) to encompass contribution of multisensory processes. For example, naturalistic studies should compare unisensory and multisensory stimulus/material formats, to measure/estimate the contribution of multisensory-driven bottom-up salience to the processes of interest. More generally, our results highlight that hypotheses about how neurocognitive functions operate in everyday situations can be built already in the laboratory, if one manipulates systematically, together and across the senses, goals, salience, and context (van Atteveldt et al. 2018; Matusz et al. 2019c). Such a cyclical approach (Matusz et al. 2019a; see also Naumann et al. 2020 for a new tool to measure ecological validity of a study) involving testing of hypotheses across laboratory and veridical real-world settings could be highly promising for successfully bridging the two, typically separately pursued types of research. As a result, such an approach could create more complete theories of naturalistic attentional control.

## Author contributions

Nora Turoman: Investigation, Formal analysis, Data curation, Software, Visualisation, Writing - original draft, Writing - review & editing.

Ruxandra I. Tivadar: Software, Writing - review & editing.

Chrysa Retsa: Software, Writing - review & editing.

Pawel J. Matusz: Conceptualization, Funding acquisition, Methodology, Resources, Formal analysis, Software, Supervision, Writing - review & editing.

Micah M. Murray: Funding acquisition, Methodology, Resources, Formal analysis, Software, Supervision, Writing - review & editing.

## Acknowledgments

We thank The EEG Brain Mapping Core of the Center for Biomedical Imaging (CIBM) for providing the infrastructure. This project was supported by the Pierre Mercier Foundation to P.J.M. Financial support was likewise provided by the Swiss National Science Foundation (grants: 320030_149982 and 320030_169206 to M.M.M., PZ00P1_174150 to P.J.M., the National Centre of Competence in research project “SYNAPSY, The Synaptic Bases of Mental Disease” [project 51AU40_125759]), and grantor advised by Carigest SA (232920) to M.M.M.. P.J.M. and M.M.M. are both supported by Fondation Asile des Aveugles.

## Abbreviations

N2pc: the N2pc event-related component
EEG: Electroencephalography
ERPs: Event-Related Potentials
TAC: Task-set Contingent Attentional Capture
MSE: Multisensory Enhancement of Attentional Capture
SOMs: Supplementary Online Materials
TCCV: target-color cue visual
NCCV: nontarget-color cue visual
TCCAV: target-color cue audiovisual
NCCAV: nontarget-color cue audiovisual
rmANOVA: repeated-measures analysis of variance
GFP: Global Field Power
TAAHC: Topographic Atomize and Agglomerate Hierarchical Clustering
D_AV_: Target Difference, difference between TCCAV and NCCAV conditions
D_V_: Target Difference, difference between TCCV and NCCV conditions
DO: Distractor Onset
MR: Multisensory Relationship

## Footnotes

1 Context has been previously defined as the “immediate situation in which the brain operates… shaped by external circumstances, such as properties of sensory events, and internal factors, such as behavioural goal, motor plan, and past experiences” (van Atteveldt et al., 2014).

2 Please see Appendix 1 for the full list of abbreviations used in the manuscript.

3 Compared to the original paradigm, we made two additional changes, to enable the Task 1 to serve as an adult control study in a developmental study (Turoman et al., 2021). We reduced the number of elements in all arrays from 6 to 4, and targets were reshaped to look like diamonds rather than rectangles. Notably, despite these changes, we have replicated here the visual and multisensory attentional control effects.

4 As part of our stimulus design and alike Matusz and Eimer (2011), we manipulated a third within-task factor, i.e., whether the distractor and the upcoming target appeared in the same compared to a different location. This manipulation was necessary for us to compute behavioural attentional capture that were the bases of our complex 4-factor analyses However, to avoid confusing the reader, we have removed the descriptions of this factor from the main text and we only refer briefly to the manipulation in the *General task procedures*.

5 While filtering following epoch creation is normally discouraged (e.g., Widmann et al. 2015), control analyses we have carried out demonstrated that our filtering procedure was necessary and did not harm the data quality within our time-window of interest (for results of control analyses, see SOMs: Justification of filtering choices).

6 Any ERP results related to Distractor Onset are unlikely to be confounded by shifted baseline due to potential dominance of one ISI type (100ms, 250ms, 450ms) over others, as no such dominance was identified in a subsample of data.

## References

Alais, D., Newell, F., & Mamassian, P. (2010). Multisensory processing in review: from physiology to behaviour. Seeing and perceiving, 23(1), 3–38.

Baess, P., Horváth, J., Jacobsen, T., & Schröger, E. (2011). Selective suppression of self-initiated sounds in an auditory stream: An ERP study. Psychophysiology, 48(9), 1276–1283.

Bevilacqua, D., Davidesco, I., Wan, L., Chaloner, K., Rowland, J., Ding, M., … & Dikker, S. (2019). Brain-to-brain synchrony and learning outcomes vary by student–teacher dynamics: Evidence from a real-world classroom electroencephalography study. Journal of cognitive neuroscience, 31(3), 401–411.

Biasiucci, A., Franceschiello, B., & Murray, M. M. (2019). Electroencephalography. Current, 29(3), R80–R85.

Brunet, D., Murray, M. M., & Michel, C. M. (2011). Spatiotemporal analysis of multichannel EEG: CARTOOL. Computational intelligence and neuroscience, 2011.

Burra, N., & Kerzel, D. (2013). Attentional capture during visual search is attenuated by target predictability: Evidence from the N2pc, Pd, and topographic segmentation. Psychophysiology, 50(5), 422–430.

Cantlon, J. F. (2020). The balance of rigor and reality in developmental neuroscience. NeuroImage, 216, 116464.

Cappe, C., Thut, G., Romei, V., & Murray, M. M. (2010). Auditory–visual multisensory interactions in humans: timing, topography, directionality, and sources. Journal of Neuroscience, 30(38), 12572–12580.

Chen, Y. C., & Spence, C. (2010). When hearing the bark helps to identify the dog: Semantically-congruent sounds modulate the identification of masked pictures. Cognition, 114(3), 389–404.

Chennu, S., Noreika, V., Gueorguiev, D., Blenkmann, A., Kochen, S., Ibánez, A., … & Bekinschtein, T. A. (2013). Expectation and attention in hierarchical auditory prediction. Journal of Neuroscience, 33(27), 11194–11205.

Chun, M. M., & Jiang, Y. (1998). Contextual cueingcueing: Implicit learning and memory of visual context guides spatial attention. Cognitive psychology, 36(1), 28–71.

Clark, A. (2013). Whatever next? Predictive brains, situated agents, and the future of cognitive science. Behavioral and brain sciences, 36(3), 181–204.

Correa, Á., Lupiáñez, J., & Tudela, P. (2005). Attentional preparation based on temporal expectancy modulates processing at the perceptual level. Psychonomic bulletin & review, 12(2), 328–334.

Coull, J. T., Frith, C. D., Büchel, C., & Nobre, A. C. (2000). Orienting attention in time: behavioural and neuroanatomical distinction between exogenous and endogenous shifts. Neuropsychologia, 38(6), 808–819.

Dassanayake, T. L., Michie, P. T., & Fulham, R. (2016). Effect of temporal predictability on exogenous attentional modulation of feedforward processing in the striate cortex. International Journal of Psychophysiology, 105, 9–16.

De Meo, R., Murray, M. M., Clarke, S., & Matusz, P. J. (2015). Top-down control and early multisensory processes: chicken vs. egg. Frontiers in integrative neuroscience, 9(17), 1–6.

Dell’Acqua, R., Sessa, P., Peressotti, F., Mulatti, C., Navarrete, E., & Grainger, J. (2010). ERP evidence for ultra-fast semantic processing in the picture–word interference paradigm. Frontiers in psychology, 1, 177.

Desimone, R., & Duncan, J. (1995). Neural mechanisms of selective visual attention. Annual Review of Neuroscience, 18(1), 193–222.

Doehrmann, O., & Naumer, M. J. (2008). Semantics and the multisensory brain: How meaning modulates processes of audio-visual integration. Brain Research, 1242, 136–50. https://doi.org/10.1016/J.BRAINRES.2008.03.071

Duncan, J., & Humphreys, G. W. (1989). Visual search and stimulus similarity. Psychological Review, 96(3), 433–458.

Eickhoff, S. B., Milham, M., & Vanderwal, T. (2020). Towards clinical applications of movie fMRI. Neuroimage, 116860.

Eimer, M. (1996). The N2pc component as an indicator of attentional selectivity. Electroencephalography and Clinical Neurophysiology, 99(3), 225–234.

Eimer, M. (2014). The neural basis of attentional control in visual search. Trends in Cognitive Sciences, 18(10), 526–535.

Eimer, M., & Kiss, M. (2008). Involuntary attentional capture is determined by task set: Evidence from event-related brain potentials. Journal of cognitive neuroscience, 20(8), 1423–1433.

Eimer, M., Kiss, M., Press, C., & Sauter, D. (2009). The roles of feature-specific task set and bottom-up salience in attentional capture: An ERP study. Journal of Experimental Psychology: Human Perception and Performance, 35(5), 1316–1328.

Ernst, M. O. (2007). Learning to integrate arbitrary signals from vision and touch. Journal of Vision, 7(5), 7–7.

Fleming, J. T., Noyce, A. L., & Shinn-Cunningham, B. G. (2020). Audio-visual spatial alignment improves integration in the presence of a competing audio–visual stimulus. Neuropsychologia, 146, 107530.

Folk, C. L., Leber, A. B., & Egeth, H. E. (2002). Made you blink! Contingent attentional capture produces a spatial blink. Perception & psychophysics, 64(5), 741–753.

Folk, C. L., Remington, R. W., & Johnston, J. C. (1992). Involuntary covert orienting is contingent on attentional control settings. Journal of Experimental Psychology: Human Perception and Performance, 18(4), 1030–1044.

Gaspelin, N., & Luck, S. J. (2019). Inhibition as a potential resolution to the attentional capture debate. Current opinion in psychology, 29, 12–18.

Gazzaley, A., & Nobre, A. C. (2012). Top-down modulation: bridging selective attention and working memory. Trends in cognitive sciences, 16(2), 129–135.

Ghazanfar, A. A., Maier, J. X., Hoffman, K. L., & Logothetis, N. K. (2005). Multisensory integration of dynamic faces and voices in rhesus monkey auditory cortex. Journal of Neuroscience, 25(20), 5004–5012.

Girelli, M., & Luck, S. J. (1997). Are the same attentional mechanisms used to detect visual search targets defined by color, orientation, and motion? Journal of Cognitive Neuroscience, 9(2), 238–253.

Golumbic, E. M. Z., Poeppel, D., & Schroeder, C. E. (2012). Temporal context in speech processing and attentional stream selection: a behavioral and neural perspective. Brain and language, 122(3), 151–161.

Green, J. J., & McDonald, J. J. (2010). The role of temporal predictability in the anticipatory biasing of sensory cortex during visuospatial shifts of attention. Psychophysiology, 47(6), 1057–1065.

Grill-Spector, K., Henson, R., & Martin, A. (2006). Repetition and the brain: neural models of stimulus-specific effects. Trends in Cognitive Sciences, 10(1), 14–23.

Guthrie, D., & Buchwald, J. S. (1991). Significance testing of difference potentials. Psychophysiology, 28(2), 240–244.

Hickey, C., Di Lollo, V., & McDonald, J. J. (2008). Target and distractor processing in visual search: Decomposition of the N2pc. Visual Cognition, 16(1), 110–113.

Hickey, C., Di Lollo, V., & McDonald, J. J. (2009). Electrophysiological indices of target and distractor processing in visual search. Journal of cognitive neuroscience, 21(4), 760–775.

Holm, S. (1979). A Simple Sequentially Rejective Multiple Test Procedure. Scandinavian Journal of Statistics, 6(2), 65–70.

Hopf, J.-M., Luck, S. J., Girelli, M., Mangun, G. R., Scheich, H., & Heinze, H.-J. (2000). Neural sources of focused attention in visual search. Cerebral Cortex, 10, 1233–1241.

Huth, A. G., Lee, T., Nishimoto, S., Bilenko, N. Y., Vu, A. T., & Gallant, J. L. (2016). Decoding the semantic content of natural movies from human brain activity. Frontiers in systems neuroscience, 10, 81.

induced gamma band responses reflect cross-modal interactions in familiar object recognition. Journal of Neuroscience, 27(5), 1090–1096.

Iordanescu, L., Guzman-Martinez, E., Grabowecky, M., & Suzuki, S. (2008). Characteristic sounds facilitate visual search. Psychonomic Bulletin & Review, 15(3), 548–554.

Jiang, J., Brashier, N. M., & Egner, T. (2015). Memory meets control in hippocampal and striatal binding of stimuli, responses, and attentional control states. Journal of Neuroscience, 35, 14885–14895.

Kingstone, A., Smilek, D., Ristic, J., Kelland Friesen, C., & Eastwood, J. D. (2003). Attention, researchers! It is time to take a look at the real world. Current Directions in Psychological Science, 12(5), 176–180.

Kiss, M., Jolicoeur, P., Dell’Acqua, R., & Eimer, M. (2008a). Attentional capture by visual singletons is mediated by top-down task set: New evidence from the N2pc component. Psychophysiology, 45(6), 1013–1024.

Kiss, M., Van Velzen, J., & Eimer, M. (2008b). The N2pc component and its links to attention shifts and spatially selective visual processing. Psychophysiology, 45, 240–249.

Klaffehn, A. L., Baess, P., Kunde, W., & Pfister, R. (2019). Sensory attenuation prevails when controlling for temporal predictability of self-and externally generated tones. Neuropsychologia, 132, 107145.

Koenig, T., Stein, M., Grieder, M., & Kottlow, M. (2014). A tutorial on data-driven methods for statistically assessing ERP topographies. Brain topography, 27(1), 72–83.

Kuo, B. C., Nobre, A. C., Scerif, G., & Astle, D. E. (2016). Top-Down activation of spatiotopic sensory codes in perceptual and working memory search.Journal of cognitive neuroscience, 28(7), 996–1009.

Laurienti, P. J., Burdette, J. H., Maldjian, J. A., & Wallace, M. T. (2006). Enhanced multisensory integration in older adults. Neurobiology of aging, 27(8), 1155–1163.

Lehmann, D. (1987). “Principles of spatial analysis,” in Methods of Analysis of Brain Electrical and Magnetic Signals, A. S. Gevins and A. Remont, Eds., pp. 309–354. Elsevier: Amsterdam, The Netherlands.

Lehmann D, Skrandies W (1980): Reference-free identification of components of checkerboard evoked multichannel potential fields. Electroencephalography in Clinical Neurology, 48, 609–621.

Lehmann, D., Ozaki, H., & Pal, I. (1987). EEG alpha map series: brain micro-states by space- oriented adaptive segmentation. Electroencephalography and clinical neurophysiology, 67(3), 271–288.

Lien, M. C., Ruthruff, E., Goodin, Z., & Remington, R. W. (2008). Contingent attentional capture by top-down control settings: converging evidence from event-related potentials. Journal of Experimental Psychology: Human Perception and Performance, 34(3), 509.

Luck, S. J., Gaspelin, N., Folk, C. L., Remington, R. W., & Theeuwes, J. (2020). Progress toward resolving the attentional capture debate. Visual Cognition, 1–21. DOI: 10.1080/13506285.2020.1848949

Luck, S. J., & Hillyard, S. A. (1994a). Electrophysiological correlates of feature analysis during visual search. Psychophysiology, 31, 291–308.

Luck, S. J., & Hillyard, S. A. (1994b). Spatial filtering during visual search: Evidence from human electrophysiology. Journal of Experimental Psychology: Human Perception and Performance, 20(5), 1000–1014.

Lunn, J., Sjoblom, A., Ward, J., Soto-Faraco, S., & Forster, S. (2019). Multisensory enhancement of attention depends on whether you are already paying attention. Cognition, 187, 38–49.

Luo, H., & Poeppel, D. (2007). Phase patterns of neuronal responses reliably discriminate speech in human auditory cortex. Neuron, 54(6), 1001–1010.

Matusz, P. J., & Eimer, M. (2013). Top-down control of audiovisual search by bimodal search templates. Psychophysiology, 50(10), 996–1009.

Matusz, P. J., & Eimer, M. (2011). Multisensory enhancement of attentional capture in visual search. Psychonomic bulletin & review, 18(5), 904.

Matusz, P. J., Wallace, M. T., & Murray, M. M. (2020). Multisensory contributions to object recognition and memory across the life span. In Multisensory Perception (pp. 135–154). Academic Press.

Matusz, P. J., Dikker, S., Huth, A. G., & Perrodin, C. (2019a). Are We Ready for Real-world Neuroscience?. Journal of cognitive neuroscience, 31(3), 327.

Matusz, P. J., Turoman, N., Tivadar, R. I., Retsa, C., & Murray, M. M. (2019b). Brain and cognitive mechanisms of top-down attentional control in a multisensory world: Benefits of electrical neuroimaging. Journal of cognitive neuroscience, 31(3), 412–430.

Matusz, P. J., Merkley, R., Faure, M., & Scerif, G. (2019c). Expert attention: Attentional allocation depends on the differential development of multisensory number representations. Cognition, 186, 171–177.

Matusz, P. J., Key, A. P., Gogliotti, S., Pearson, J., Auld, M. L., Murray, M. M., & Maitre, N. L. (2018). Somatosensory plasticity in pediatric cerebral palsy following constraint- induced movement therapy. Neural Plasticity, 2018.

Matusz, P. J., Wallace, M. T., & Murray, M. M. (2017). A multisensory perspective on object memory. Neuropsychologia, 105, 243–252.

Matusz, P. J., Retsa, C., & Murray, M. M. (2016). The context-contingent nature of cross- modal activations of the visual cortex. Neuroimage, 125, 996–1004.

Matusz, P. J., Thelen, A., Amrein, S., Geiser, E., Anken, J., & Murray, M. M. (2015a). The role of auditory cortices in the retrieval of single-trial auditory-visual object memories. European Journal of Neuroscience, 41(5), 699–708.

Matusz, P. J., Broadbent, H., Ferrari, J., Forrest, B., Merkley, R., & Scerif, G. (2015b). Multimodal distraction: Insights from children’s limited attention. Cognition, 136, 156–165.

McNab, F., & Klingberg, T. (2008). Prefrontal cortex and basal ganglia control access to working memory. Nature Neuroscience, 11, 103–107.

Michel, C. M., & Murray, M. M. (2012). Towards the utilization of EEG as a brain imaging tool. Neuroimage, 61(2), 371–385.

Miniussi C, Wilding EL, Coull JT, Nobre AC. (1999). Orienting atten- tion in the time domain: modulation of potentials. Brain, 122, 1507–18.

Murray, M. M., Brunet, D., & Michel, C. M. (2008). Topographic ERP analyses: a step-by-step tutorial review. Brain topography, 20(4), 249–264.

Murray, M. M., Thelen, A., Thut, G., Romei, V., Martuzzi, R., & Matusz, P. J. (2016a). The multisensory function of the human primary visual cortex. Neuropsychologia, 83, 161–169.

Murray, M. M., Lewkowicz, D. J., Amedi, A., & Wallace, M. T. (2016b). Multisensory processes: a balancing act across the lifespan. Trends in Neurosciences, 39(8), 567–579.

Murray, M. M., Michel, C. M., De Peralta, R. G., Ortigue, S., Brunet, D., Andino, S. G., & Schnider, A. (2004). Rapid discrimination of visual and multisensory memories revealed by electrical neuroimaging. Neuroimage, 21(1), 125–135.

Naccache, L., Blandin, E., & Dehaene, S. (2002). Unconscious masked priming depends on temporal attention. Psychological Science, 13(5), 416–424.

Nastase, S. A., Goldstein, A., & Hasson, U. (2020). Keep it real: rethinking the primacy of experimental control in cognitive neuroscience. Neuroimage. In print.

Naumann, S., Byrne, M. L., de la Fuente, L. A., Harrewijn, A., Nugiel, T., Rosen, M. L., … & Matusz, P. J. (2020). Assessing the degree of ecological validity of your study: Introducing the Ecological Validity Assessment (EVA) Tool. PsyArXiv. DOI: 10.3l234/osf.io/qb9tz.

Neel, M. L., Yoder, P., Matusz, P. J., Murray, M. M., Miller, A., Burkhardt, S., … & Maitre, N. L. (2019). Randomized controlled trial protocol to improve multisensory neural processing, language and motor outcomes in preterm infants. BMC Pediatrics, 19(1), 1–10.

Noonan, M. P., Crittenden, B. M., Jensen, O., & Stokes, M. G. (2018). Selective inhibition of distracting input. Behavioural brain research, 355, 36–47.

Peelen, M. V., & Kastner, S. (2014). Attention in the real world: toward understanding its neural basis. Trends in cognitive sciences, 18(5), 242–250.

Perrin, F., Pernier, J., Bertnard, O., Giard, M. H., & Echallier, J. F. (1987). Mapping of scalp potentials by surface spline interpolation. Electroencephalography and clinical neurophysiology, 66(1), 75–81.

Press, C., Kok, P., & Yon, D. (2020). The perceptual prediction paradox. Trends in Cognitive Sciences, 24(1), 13–24.

Raij, T., Ahveninen, J., Lin, F. H., Witzel, T., Jääskeläinen, I. P., Letham, B., … & Hämäläinen, M. (2010). Onset timing of cross-sensory activations and multisensory interactions in auditory and visual sensory cortices. European Journal of Neuroscience, 31(10), 1772–1782.

Raij, T., Uutela, K., & Hari, R. (2000). Audiovisual integration of letters in the human brain. Neuron, 28(2), 617–625.

Retsa, C., Matusz, P. J., Schnupp, J. W., & Murray, M. M. (2018). What’s what in auditory cortices?. NeuroImage, 176, 29–40.

Retsa, C., Matusz, P. J., Schnupp, J. W., & Murray, M. M. (2020). Selective attention to sound features mediates cross-modal activation of visual cortices. Neuropsychologia, 144, 107498.

Richter, D., Ekman, M., & de Lange, F. P. (2018). Suppressed sensory response to predictable object stimuli throughout the ventral visual stream. Journal of Neuroscience, 38(34), 7452–7461.

Rohenkohl, G., Gould, I. C., Pessoa, J., & Nobre, A. C. (2014). Combining spatial and temporal expectations to improve visual perception. Journal of vision, 14(4), 8–8.

Sadeh, T., Shohamy, D., Levy, D. R., Reggev, N., & Maril, A. (2011). Cooperation between the hippocampus and the striatum during episodic encoding. Journal of Cognitive Neuroscience, 23(7), 1597–1608.

Sarmiento, B. R., Matusz, P. J., Sanabria, D., & Murray, M. M. (2016). Contextual factors multiplex to control multisensory processes. Human brain mapping, 37(1), 273–288.

Sawaki, R., & Luck, S. J. (2010). Capture versus suppression of attention by salient singletons: Electrophysiological evidence for an automatic attend-to-me signal. Attention, Perception, & Psychophysics, 72(6), 1455–1470.

Schröger, E., Marzecová, A., & SanMiguel, I. (2015). Attention and prediction in human audition: A lesson from cognitive psychophysiology. European Journal of Neuroscience, 41(5), 641–664.

Shamay-Tsoory, S. G., & Mendelsohn, A. (2019). Real-life neuroscience: an ecological approach to brain and behavior research. Perspectives on Psychological Science, 14(5), 841–859.

Soto-Faraco, S., Kvasova, D., Biau, E., Ikumi, N., Ruzzoli, M., Morís-Fernández, L., & Torralba, M. (2019). Multisensory interactions in the real world. Cambridge University Press.

Southwell, R., Baumann, A., Gal, C., Barascud, N., Friston, K., & Chait, M. (2017). Is predictability salient? A study of attentional capture by auditory patterns. Philosophical Transactions of the Royal Society B: Biological Sciences, 372(1714), 20160105.

Spierer, L., Manuel, A. L., Bueti, D., & Murray, M. M. (2013). Contributions of pitch and bandwidth to sound-induced enhancement of visual cortex excitability in humans. Cortex, 49(10), 2728–2734.

Sui, J., He, X., & Humphreys, G. W. (2012). Perceptual effects of social salience: evidence from self-prioritization effects on perceptual matching. Journal of Experimental Psychology: Human perception and performance, 38(5), 1105.

Summerfield, C., & Egner, T. (2009). Expectation (and attention) in visual cognition. Trends in cognitive sciences, 13(9), 403–409.

Summerfield, J. J., Lepsien, J., Gitelman, D. R., Mesulam, M. M., & Nobre, A. C. (2006). Orienting attention based on long-term memory experience. Neuron, 49(6), 905–916.

Sun, Y., Fuentes, L. J., Humphreys, G. W., & Sui, J. (2016). Try to see it my way: Embodied perspective enhances self and friend-biases in perceptual matching. Cognition, 153, 108–117.

Talsma, D., & Woldorff, M. G. (2005). Selective attention and multisensory integration: multiple phases of effects on the evoked brain activity. Journal of Cognitive Neuroscience, 17,1098–1114.

Ten Oever, S., & Sack, A. T. (2015). Oscillatory phase shapes syllable perception. Proceedings of the National Academy of Sciences, 112(52), 15833–15837.

Ten Oever, S., Romei, V., van Atteveldt, N., Soto-Faraco, S., Murray, M. M., & Matusz, P. J. (2016). The COGs (context, object, and goals) in multisensory processing. Experimental brain research, 234(5), 1307–1323.

Theeuwes, J. (1991). Cross-dimensional perceptual selectivity. Perception & Psychophysics, 50(2), 184–193.

Thelen, A., Talsma, D., & Murray, M.M. (2015). Single-trial multisensory memories affect later auditory and visual object discrimination. Cognition 138, 148–160.

Thorpe, S., Fize, D., & Marlot, C. (1996). Speed of processing in the human visual system. Nature, 381(6582), 520–522.

Tivadar, R. I., & Murray, M. M. (2019). A primer on electroencephalography and event- related potentials for organizational neuroscience. Organizational Research Methods, 22(1), 69–94.

Tivadar, R. I., Knight, R. T., & Tzovara, A. (2021). Automatic Sensory Predictions: A Review of Predictive Mechanisms in the Brain and Their Link to Conscious Processing. Frontiers in Human Neuroscience, 438.

Tovar, D. A., Murray, M. M., & Wallace, M. T. (2020). Selective enhancement of object representations through multisensory integration. Journal of Neuroscience. In press. DOI: https://doi.org/10.1523/JNEUROSCI.2139-19.2020

Treisman, A. M., & Gelade, G. (1980). A feature-integration theory of attention. Cognitive Psychology, 12(1), 97–136.

Turoman, N., Tivadar, R. I., Retsa, C., Maillard, A. M., Scerif, G., and Matusz, P. J. (2021a). The development of attentional control mechanisms in multisensory environments. Developmental Cognitive Neuroscience, 48, 100930.

Turoman, N., Tivadar, R. I., Retsa, C., Maillard, A. M., Scerif, G., and Matusz, P. (2021b). Uncovering the mechanisms of real-world attentional control over the course of primary education. Mind, Brain, & Education. In press.

Tzovara, A., Murray, M. M., Michel, C. M., & De Lucia, M. (2012). A tutorial review of electrical neuroimaging from group-average to single-trial event-related potentials. Developmental neuropsychology, 37(6), 518–544.

Van Atteveldt, N., Murray, M. M., Thut, G., & Schroeder, C. E. (2014). Multisensory integration: flexible use of general operations. Neuron, 81(6), 1240–1253.

van Atteveldt, N., van Kesteren, M. T. R., Braams, B.; Krabbendam, L. (2018). Neuroimaging of learning and development: improving ecological validity. Frontline Learning Research, 6 (3), 186–203. DOI: 10.14786/flr.v6i3.366.

Van der Burg, E., Talsma, D., Olivers, C. N. L., Hickey, C., & Theeuwes, J. (2011). Early multisensory interactions affect the competition among multiple visual objects. NeuroImage, 55, 1208–1218.

van Moorselaar, D., & Slagter, H. A. (2019). Learning what is irrelevant or relevant: Expectations facilitate distractor inhibition and target facilitation through distinct neural mechanisms. Journal of Neuroscience, 39(35), 6953–6967.

van Moorselaar, D., & Slagter, H. A. (2020). Inhibition in selective attention. Annals of the New York Academy of Sciences, 1464(1), 204.

van Moorselaar, D., Daneshtalab, N., & Slagter, H. (2020). Neural mechanisms underlying distractor inhibition on the basis of feature and/or spatial expectations. bioRxiv.

Vanderwal, T., Eilbott, J.; Castellanos, F. X (2019). Movies in the magnet: Naturalistic paradigms in developmental functional neuroimaging. Developmental Cognitive Neuroscience, 36, 100600.

Vaughan Jr, H. G. (1982). The neural origins of human event-related potentials. Annals of the New York Academy of Sciences, 388(1), 125–138.

Widmann, A., Schröger, E., & Maess, B. (2015). Digital filter design for electrophysiological data–a practical approach. Journal of Neuroscience Methods, 250, 34–46.

Wu, R., Nako, R., Band, J., Pizzuto, J., Ghoreishi, Y., Scerif, G., & Aslin, R. (2015). Rapid attentional selection of non-native stimuli despite perceptual narrowing. Journal of Cognitive Neuroscience, 27(11), 2299–2307.

Yuval-Greenberg, S., & Deouell, L. Y. (2007). What you see is not (always) what you hear: induced gamma band responses reflect cross-modal interactions in familiar object recognition. Journal of Neuroscience, 27(5), 1090–1096.

