## Supplementary Online Materials for "Towards understanding how we pay attention in naturalistic visual search settings" (from another version of this preprint: version 3, 2021-07-12)

**Supplementary Online Materials (SOMs)**

### Supplementary Materials & Methods

#### Power and effect size calculations

Previous studies with comparable experimental tasks normally involved smaller samples (Eimer, Kiss, Press & Sauter, 2009; Matusz & Eimer, 2011). In the original study that served as a basis for the current study (Matusz & Eimer, 2011, Task 2), the closest to our TAC and MSE effects in behaviour were their Cue Colour x Spatial Cueing interaction and Tone Presence x Spatial Cueing interaction. In that study, the power of the Cue Colour x Spatial Cueing interaction was 0.99 with a sample size of 22 and an effect size of 0.6, while the power of the Tone Presence x Spatial Cueing interaction was 0.92 with a sample size of 22 and an effect size of 0.1. A post-hoc power analysis on our present data revealed sufficient statistical power for testing our hypothesised TAC effects (0.99 for a main effect of Cue Colour size of 39, effect size of 0.9) and MSE effects (0.99 for a main effect of Cue Modality, sample size of 39, effect size of 0.2) in behaviour. For ERP analyses, we could only estimate the power and effect size necessary to detect TAC effects in canonical N2pc analyses. Here, the closest compatible effect was the Cue Condition x Contralaterality interaction from Eimer et al. (2009). This interaction reached a power of 0.99 with a sample size of 12 and an effect size of 0.6. We, again, conducted a post-hoc power analysis on our N2pc data from Task 2 and found that there was sufficient statistical power to detect TAC in N2pc (0.99 for a main effect of Cue Colour size of 39, effect size of 0.4). The power of MSE and contextual effects in N2pc could not be estimated as no previous studies have investigated such effects in distractor-locked N2pc's.

#### Control Inter-stimulus interval (ISI) analyses

It was possible that the distractor-elicited ERPs in our Tasks with variable interstimulus intervals (ISI's) had shifted ERP baselines, which would act as a confounding influence on effects related to the factor of Distractor Onset (Distractor Onset). In order to investigate this possibility, we compared the proportion of trials for each ISI type (short [100ms] vs. medium [250ms] vs. long [450ms]) in a subset of our data to see if there indeed was an imbalance that could have driven our effects. Here, only data from Tasks 1 and 3 could be included, as these were the Tasks where ISI duration varied (in Exps. 2 and 4 the duration was always 450ms). More broadly, as the ISI duration was programmed to vary randomly for each subject, we can safely assume that the frequency of ISI's in the subset of 11 participants generalises to the entire dataset. We compared the numbers of trials on behavioural datasets across the different ISIs, as the EEG output files did not contain any information about ISI duration on a trial-by-trial basis. That said, we knew that our ERP analyses did not include trials with inaccurate and missed responses, which were also "filtered out" during our initial cleaning of the behavioural data. With that, we present the investigation of the clean behavioural datasets below.

While the numbers of trials per ISI type were numerically unequal, that was to be expected, as the total number of trials in each Task was 512, which is not divisible by the number of ISI types. That is, a perfectly balanced number of trials per ISI type would be impossible in practice, with 170.67 trials per ISI type. To test if this inequality in trial numbers was statistically significant, and would thus warrant concern about confounding influences on Distractor Onset-related effects in ERPs, we submitted the numbers of trials to a set of one-way ANOVAs. First, we analysed the numbers of trials in Task 1 and 3 combined and found that the numbers of trials were not significantly different ($F_{(2, 42)}$ = 0.8, $p$ = 0.4, $\eta_p^2$ = 0.3). Next, we looked at each Task individually. Again, we found that the numbers of trials were not significantly different for Task 1 ($F_{(2, 20)}$ = 0.9, $p$ = 0.4, $\eta_p^2$ = 0.1) or Task 3 ($F_{(2, 20)}$ = 0.2, $p$ = 0.9, $\eta_p^2$ = 0.01). With this, we concluded that there was no disproportionate involvement of any one of the 3 ISI types that could skew the results.

#### Justification of filtering choices

##### 1. Bandpass and first Notch

In Supplemental Figure 1, on the far left, top to bottom panels, we plotted the raw signal, the 0.1-40hHz band-passed signal, and the 0.1-40Hz band-passed + 50Hz Notched signal, respectively, in teal to signify the time domain. The incremental reduction in noise after each step is readily observable. It is also clear that the Notch filter does dramatically reduce noise, even though, technically, that Notch should be encompassed in the initial band-pass filter. In the next column of images, to the right, we plotted the same sequence of signals but in purple, to signify the frequency domain. From the top panel, it is clear that the raw data had a large amount of 50Hz noise, as well as some noise from other frequencies as well. The band-pass filter attenuated, but certainly did not remove, the 50Hz noise, whereas most of the other noise was removed. Notably, after the additional Notch, the 50Hz noise was completely removed. In the next two columns to the right, we plotted difference plots showing the difference in the signal (teal) and frequency spectrum (purple) at each step of the filtering process. From these plots, it is clear that at each of the two steps, there is a non-negligible amount of noise that was removed. We thus conclude that the band-pass filter was demonstrably imperfect, but that the additional Notch was necessary to fully remove noise.

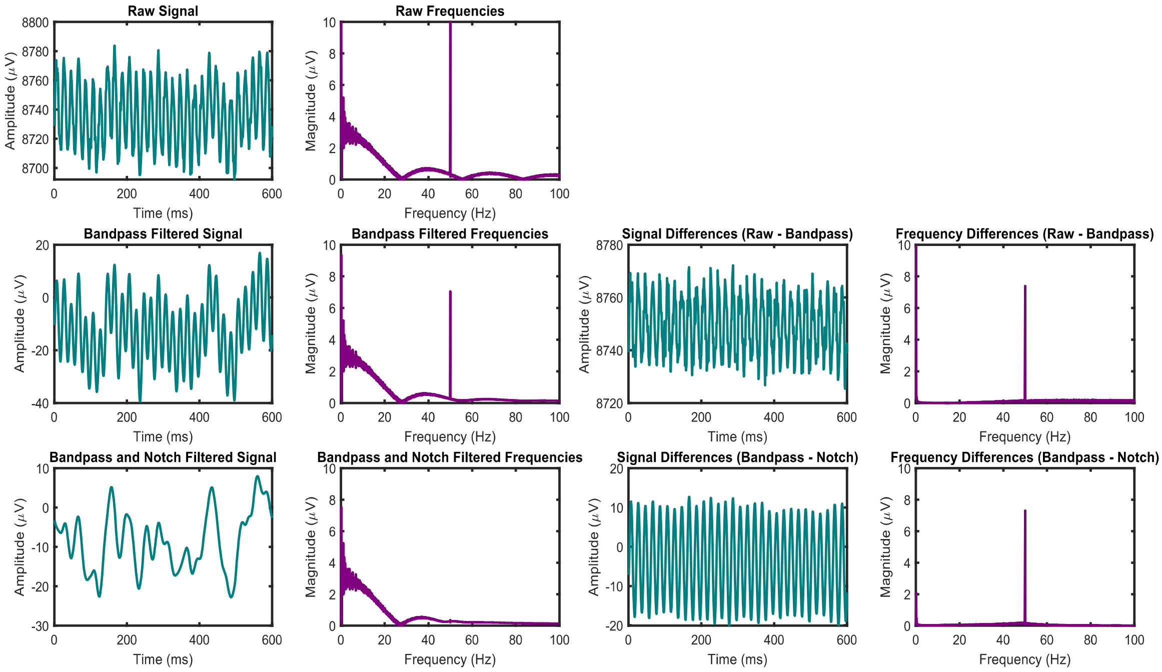

**Supplemental Figure 1.** The raw signal, the 0.1-40hHz band-passed signal, and the 0.1-40Hz band-passed + 50Hz Notched signal, respectively (in teal to signify the time domain; far left, top to bottom panels). In the column of images to the right, the same sequence of signals plotted (in purple, to signify the frequency domain).

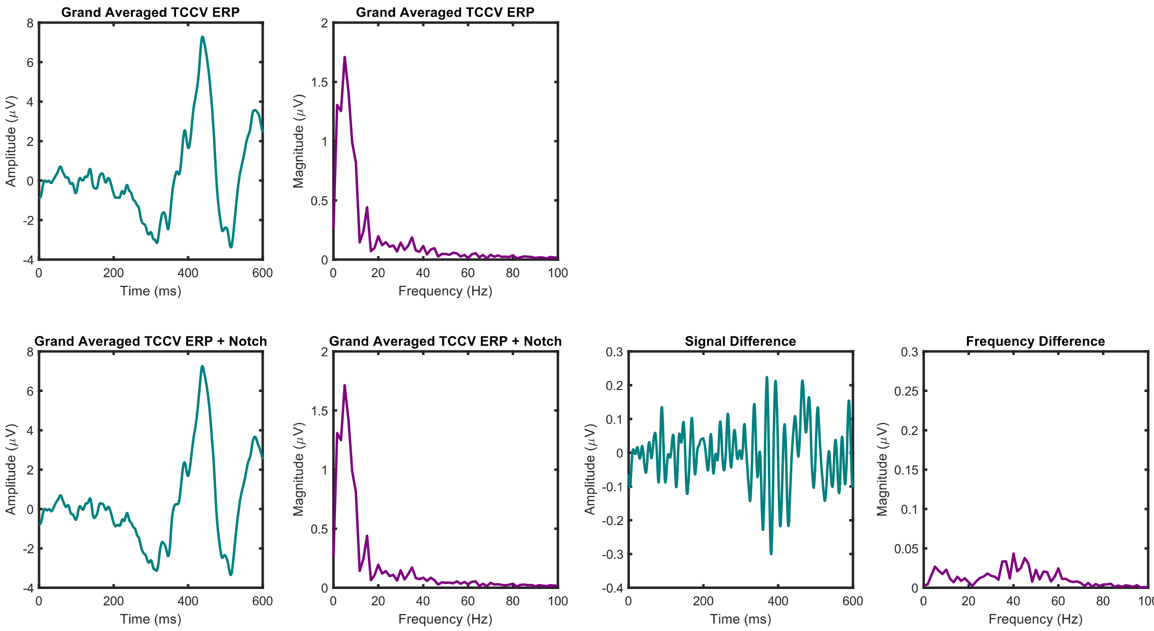

**Supplemental Figure 2.** Grand-averaged TCCV ERP from the same participant, block, and Task as in the previous step, before the second 50Hz notch (upper row), and after the second 50Hz notch (lower row) (in teal to signify the time domain; far left, top to bottom panels). In the column of images to the right, the same sequence of signals plotted (in purple, to signify the frequency domain).

#### 2. Second Notch after epoch creation

In Supplemental Figure 2, on the far left, top to bottom panels, we plotted the grand-averaged TCCV ERP from the same participant, block, and Task as in the previous step, before the second 50Hz notch (upper row), and after the second 50Hz notch (lower row). Here we see that the shape of the signal was not dramatically altered by the addition of the second Notch, and that the Notch had only slight effects (e.g. the edges on the target P1 and N2pc-like peak are somewhat smoother). In the next column of images to the right we plotted the frequency spectrum, where differences are even harder to notice. However, in the difference plots in the next columns to the right, it appears that noise ~50Hz was eliminated following the application of this Notch.

While filtering after grand-averaging should generally be discouraged, our analysis shows that, in the time-window of interest (until 300ms post-distractor), the second notch did not distort the signal to unrepresentative levels (the shapes of the peaks and troughs of interest were preserved), but it did remove noise that survived the first filtering stage, which was its intended purpose.

#### 3. Dirac function test

Finally, in order to make the filter characteristics as explicit as possible, we applied our Bandpass filter (0.1 - 40Hz) and Notch (50Hz) onto a Dirac function. Supplemental Figure 3 shows the impulse responses of the bandpass and notch (teal, left column) and frequency responses of the bandpass and notch (purple, right column). From these plots, we can see the shape of the filters, and that, for our bandpass filter, frequencies from 0.1 and above pass with a good cutoff.

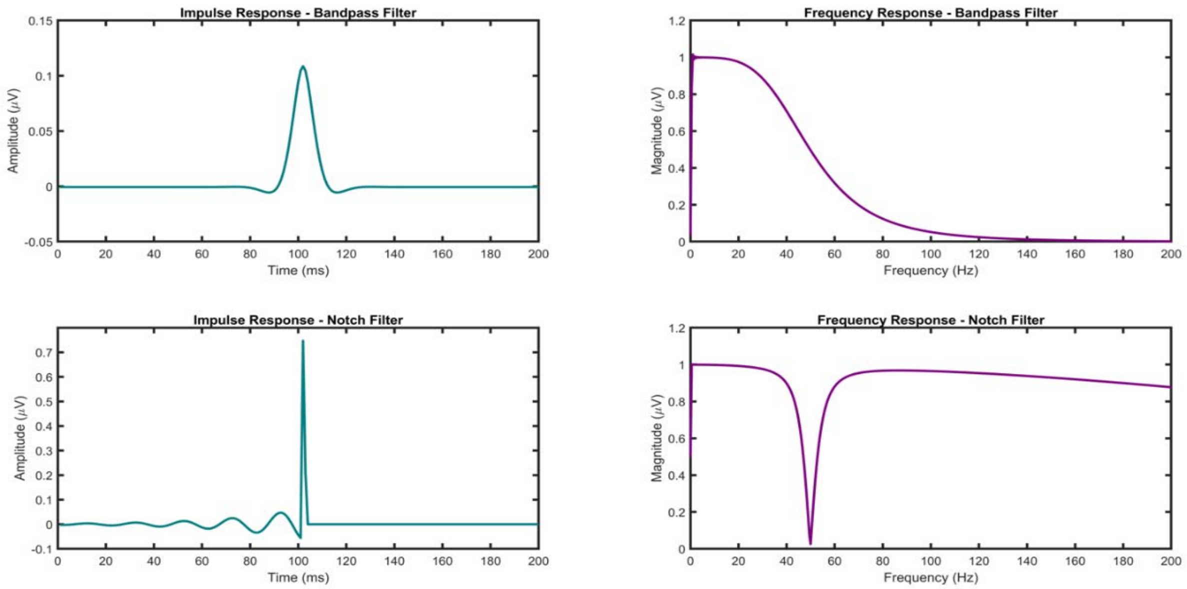

**Supplemental Figure 3.** The impulse responses of the bandpass and notch (teal, left column) and frequency responses of the bandpass and notch (purple, right column), following applying onto a Dirac function the Bandpass filter (0.1 - 40Hz) and Notch (50Hz) filter that we used for our EEG/ERP analyses.

### Supplemental Behavioural Results

#### Hypothesis-driven analyses: Interactions between TAC and MSE and contextual factors

#### RT results

*Omnibus 2 × 2 × 2 × 2 rmANOVA: Full Results*

There were no significant main effects of the two contextual factors (Multisensory Relationship, $F_{(1, 38)} = 1.9$, $p = 0.18$; Distractor Onset, $F_{(1, 38)} = 0.3$, $p = 0.6$). No other interactions than those described in the main text were statistically reliable (all $F$'s < 3, $p$'s > 0.1).

The follow-up analysis of the 2-way Distractor Colour x Multisensory Relationship interaction as a function of Multisensory Relationship showed that TCC distractors were found to elicit larger behavioural capture effects than NCC distractors, both for Arbitrary (TCCs vs. NCCs: 45ms vs. -2ms, $t_{(38)} = 32.5$, $p < 0.001$) and Congruent (TCCs vs. NCCs: 40ms vs. -0.2ms, $t_{(38)} = 25.6$, $p < 0.001$) distractors.

#### Error rates

Error data were not normally distributed, and thus a 1-way Kruskal–Wallis $H$ test was conducted between Tasks, and Durbin tests were conducted for each Task separately. Overall, error rates differed significantly between Tasks, $\chi^2(3) = 34.7$, $p < 0.001$. Error rates were highest in Task 2 (6.1%), followed by Task 1 (5.8%), both of which involved a purely arbitrary presentation of the visual and auditory attributes of multisensory distractors, i.e., the Arbitrary condition of Multisensory Relationship. Next were Task 3 (2.6%) and Task 4 (2.5%), where the auditory and visual attributes were semantically congruent. Across Tasks, Durbin tests revealed that error rates were modulated by Distractor-Target Location, but not by Distractor Colour or Distractor Modality ($p$'s > 0.1). Thus, in Task 1, $\chi^2(1) = 6.9$, $p = 0.009$, fewer errors were made on trials where the distractor and target location were the same (5.3%) than when they were different (6.3%). In Task 2, $\chi^2(1) = 13.9$, $p < 0.001$, again, fewer errors were made on trials where the distractor and target location were the same (5.4%) than when they were different (6.8%). In Task 3, $\chi^2(1) = 10.4$, $p < 0.001$, as well, fewer errors were made when the distractor and target location were the same (2.3%) than when they were different (3%). Finally, in Task 4, $\chi^2(1) = 8.3$, $p = 0.004$, fewer errors were made when the distractor and target location were the same (2.3%) than when they were different (2.9%).

### Supplemental ERP & Electrical Neuroimaging Results

#### Lateralised ERP analyses

##### Interaction of TAC with contextual factors

*Canonical N2pc analysis.* The 2 × 2 × 2 × 2 rmANOVA on the lateralised distractor-elicited ERPs revealed a main effect of Distractor Colour, $F_{(1, 38)} = 5.5$, $p = 0.03$, $\eta_p^2 = 0.1$, thus confirming the presence of TAC in the N2pc analysis. This effect was driven by a stronger mean amplitude of contra-ipsi difference for TCCs (-0.08µV) than for NCCs (-0.02µV; Main text **Error! Reference source not found.**).

*Lateralised electrical neuroimaging analyses of the N2pc component.* We investigated whether distractor-elicited Task-set contingent attentional capture-related modulations of lateralised ERPs by contextual factors resulted from response strength differences. We did so by conducting a 2 × 2 × 2 × 2 rmANOVA on average Global field power values over the N2pc time-window. Also, to investigate whether network-based differences contributed to modulations of distractor-elicited lateralised ERPs by contextual factors, we conducted a segmentation over the entire post-distractor time-period. The segmentation revealed 13 clusters which explained 75.4% of GEV in the group-averaged difference ERPs. Over the N2pc time-window (180-300ms), there were 3 template maps (hereafter Map1, Map2, and Map3), the presence of which over the N2pc time-window was submitted to a 2 × 2 × 2 × 2 x 3 rmANOVA.

GFP measured on the lateralised ERPs showed task-set contingent attentional capture, as evidenced by a main effect of Distractor Colour, $F(1, 38) = 10.2$, $p = 0.003$, $\eta p^2 = 0.2$, such that GFP was larger for TCC distractors (0.6µV) than for NCC distractors (0.5µV).

There was also evidence for TAC in the lateralised ERP topography, as the main effect of Map, $F_{(1.6, 61.4)} = 54.7$, $p < 0.001$, $\eta_p^2 = 0.6$, was modulated by Distractor Colour as shown by a 2-way Map x Distractor Colour interaction, $F_{(1.5, 58.4)} = 5.4$, $p = 0.02$, $\eta_p^2 = 0.1$. A follow-up of this interaction by Distractor Colour revealed that Map2 was present longer for TCC (25ms) than NCC distractors (14ms), $t_{(38)} = 3$, $p = 0.01$, and Map1 was present shorter for TCC (40ms) than NCC distractors (52ms), $t_{(38)} = 3.1$, $p = 0.01$. Meanwhile, Map3 was not differently present between TCC (56ms) and NCC distractors (56ms, $t_{(38)} = 0.1$, $p = 1$ (Hereafter, map duration differences that were not reported here were not statistically significant, $p$'s > 0.1). These results suggest that TAC is driven by a set of network-based mechanisms, i.e., the processing of TCC distractors by Map2, and the processing of NCC distractors by Map1.

A follow-up of the 2-way Map x Distractor Colour interaction ($F_{(1.5, 58.4)} = 5.4$, $p = 0.02$, $\eta_p^2 = 0.1$) by Map revealed that Map2 was present shorter than other maps for both TCC and NCC distractors. Specifically, for TCC distractors, Map2 (25ms) was shorter than Map1 (41ms), $t_{(38)} = 3.3$, $p = 0.008$, and than Map3 (56ms), $t = 3.2$, $p < 0.001$. Similarly, for NCC distractors, Map2 (14ms) was also shorter than Map1 (52ms), $t_{(38)} = 5$, $p < 0.001$, and than Map3 (56ms), $t_{(38)} = 8.7$, $p < 0.001$. A follow-up analysis of the Distractor Onset x Map interaction ($F_{(1.8, 67.7)} = 5.9$, $p = 0.006$, $\eta_p^2 = 0.1$) revealed that Map2 was shorter than the other two maps both for Predictable and Unpredictable distractors. Specifically, for Unpredictable distractors, Map2 (22ms) was shorter than Map1 (48ms), $t_{(38)} = 6.3$, $p < 0.001$, and than Map3 (52ms), $t = 7.4$, $p < 0.001$. Meanwhile for Predictable distractors, Map2 (17ms) was also shorter than Map1 (45ms), $t_{(38)} = 6.8$, $p < 0.001$, and than Map3 (60ms), $t_{(38)} = 10.4$, $p < 0.001$.

##### Interaction of MSE with contextual factors

*Canonical N2pc analysis.* The 2 × 2 × 2 × 2 rmANOVA on the lateralised distractor-elicited ERPs revealed no significant main effect of Distractor Modality, thus providing no evidence for MSE in the traditional N2pc analyses. No other main effects or interactions in the canonical N2pc analysis were significant, including the main effects of Multisensory Relationship or Distractor Onset, $F$'s < 3, $p$'s > 0.08, or any interactions between these two factors and any other factors (all $F$'s < 1).

*Lateralised Electrical Neuroimaging analyses of the N2pc component.* There was no evidence for MSE in GFP, $F_{(1, 38)} = 0.1$, $p = 0.8$, or in topography, $F_{(1.7, 63.9)} = 0.7$, $p = 0.5$. MSE was also not found to be modulated by either one of the contextual factors in N2pc, GFP, or topography (all $F$'s < 1).

The behaviourally-observed interaction between TAC and MSE was not found in N2pc ($F_{(1, 38)} = 2.2$, $p = 0.2$), GFP, or topography ($F$'s < 1, $p$'s > 0.1).

#### Nonlateralised ERP analyses

##### Interaction of Target Difference with contextual factors.

*Electrode-wise voltage analyses.* The ANOVA revealed a main effect of Target Difference, at 53 – 99ms and 141 – 179ms, where voltages were larger for $D_{AV}$ (TCCAV – NCCAV difference) than for $D_V$ (TCCV – NCCV difference) at both time-periods (all $p$'s < 0.05). In terms of contextual factors, there was also a significant main effect of Multisensory Relationship at 140 – 162ms, such that voltage amplitudes were higher for Congruent than for Arbitrary distractors (all $p$'s < 0.05). Target Difference and Multisensory Relationship interacted with each other, at the following timepoints: 65 – 103ms, 143 – 171ms, and 194 – 221ms (all $p$'s < 0.05). The interaction was followed up as a function of Multisensory Relationship. For Congruent distractors, significantly larger ERP amplitudes were observed for $D_{AV}$ than for $D_V$ at 48 – 97ms, 137 – 182ms, and 191 – 231ms (all $p$'s < 0.05; Main text **Error! Reference source not found.**C). In contrast, for Arbitrary distractors, there were no significant differences between the levels of Target Difference.

There was no significant main effect of Distractor Onset, as the data did not meet the temporal criterion at any timepoint. Other interactions failed to meet either the spatial or temporal criterion for significance, or both.

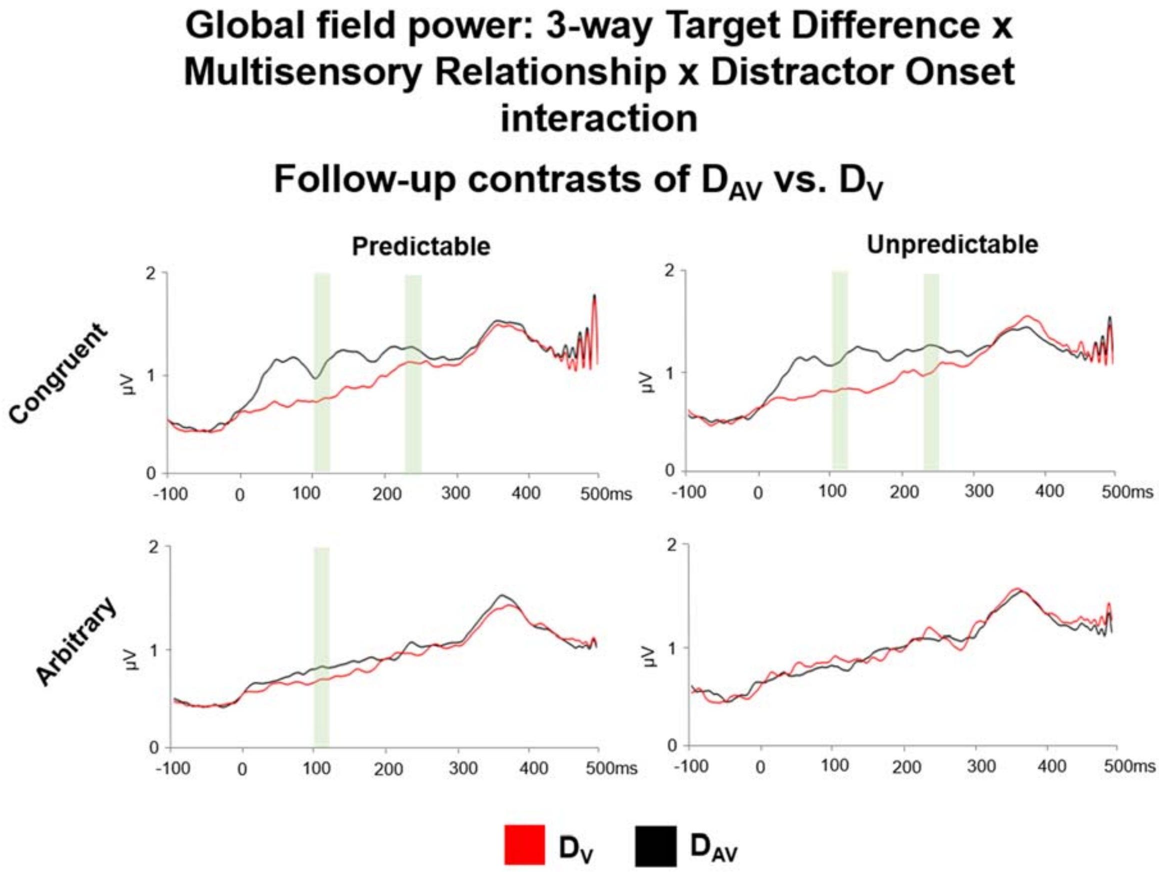

**Supplemental Figure 4.** Results of 3-way Target Difference x Multisensory Relationship x Distractor Onset interaction post-hoc tests in GFP across $D_V$ and $D_{AV}$ difference within each of the 4 context conditions across the whole post-distractor time-window. The shaded areas indicate time-windows where the difference was statistically significant.

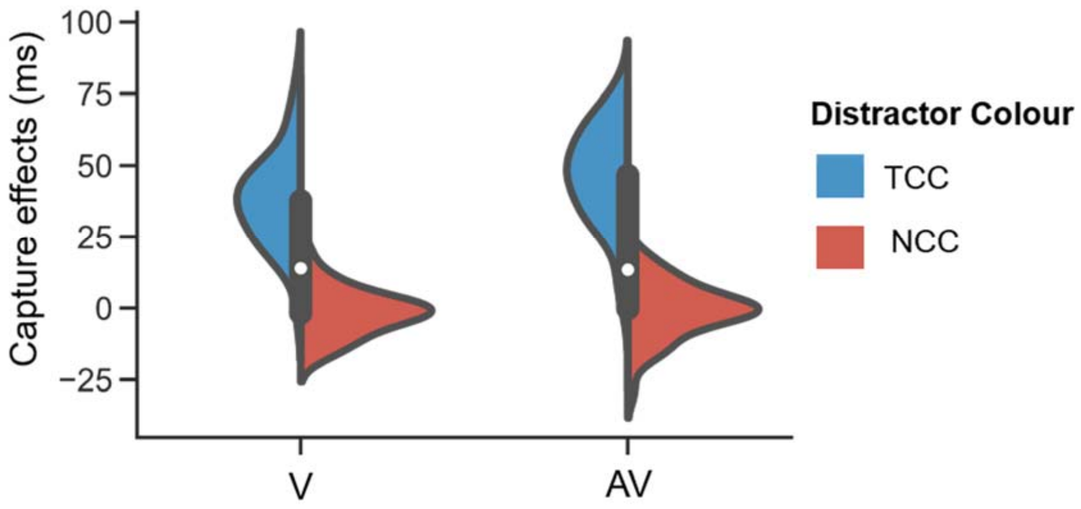

**Supplemental Figure 5.** Overall behavioural capture effects elicited by target-colour distractors (TCC) were larger than that of non-target colour (NCC) distractors, overall, confirming task-set contingent attentional capture. Likewise, there was MSE, as in the original Matusz and Eimer (2011) study. However, when the data from all 4 Tasks were treated together, a 2-way Distractor Colour x Distractor Modality interaction demonstrated that MSE was limited to TCC distractors.

*GFP analyses.* The 2 x 2 x 2 ANOVA was conducted on GFP values over the entire post-distractor period in a millisecond-by-millisecond fashion. There was a significant main effect of Multisensory

Relationship at the following timeframes: 23 – 180ms, 188 – 234ms, 242 – 261ms, where GFP was larger for Arbitrary than Congruent distractors across all time-periods (all p's < 0.05). The main effect of Distractor Onset was also significant, at the following timeframes: 13 – 34ms, 97 – 118ms, 304 – 335ms, where GFP was larger for Unpredictable than Predictable distractors for all but the middle time-period, where Predictable distractors had a larger GFP than Unpredictable distractors (all p's < 0.05)

We explored the 3-way Target Difference × Multisensory Relationship × Distractor Onset interaction first by testing the statistical significance of the $D_V$ and $D_{AV}$ difference within each of the 4 context conditions (i.e., Arbitrary Unpredictable, Arbitrary Predictable, Congruent Unpredictable, and Congruent Predictable) across the whole post-distractor and pre-target time-window. These differences were reliable for all conditions besides the Unpredictable Arbitrary condition (see Supplemental Figure 4). Second, we explored the 3-way interaction to assess the strength of task-set contingent attentional capture-related responses across the 4 contexts by comparing just the $D_V$ responses. For Congruent distractors, $D_V$ GFPs were not significantly different across Predictable and Unpredictable distractors (the difference lasted across 234-245ms and so did not meet the temporal criterion within the two time-windows of the 3-way interaction). Another post-hoc test that did not fully meet the temporal criterion was a trend for larger GFPs for Unpredictable Arbitrary than Unpredictable Congruent distractors. We also explored the 3-way interaction to compare the AV-induced enhancements between $D_{AV}$ and $D_V$ across the 4 contexts, by directly comparing the $D_{AV} – D_V$ subtractions. The AV-enhancements for Predictable Arbitrary compared to Unpredictable Arbitrary in both time-windows that did not meet the temporal criteria spanned 102-112ms and 242-249ms post-distractor.

In GFP, the Target Differences interacted separately with both contextual factors. First, the 2-way Target Difference x Multisensory Relationship interaction spanned a large part of the post-distractor time period (23 – 255ms). We followed up the interaction by Multisensory Relationship, as the interest was to investigate how contextual factors modulate Target Difference. For Congruent distractors, significantly larger GFP amplitudes were observed for $D_{AV}$ than for $D_V$ over 11 – 255ms (p < 0.05). In contrast, for Arbitrary distractors, no significant differences between the levels of Target Difference were observed. Second, the Target Difference x Distractor Onset interaction was significant at 88 – 127ms, and we followed it up as a function of Distractor Onset. For both Unpredictable and Predictable distractors, significantly larger GFP amplitudes were observed for $D_{AV}$ than for $D_{V,}$ but at different latencies (Unpredictable: 8 – 102ms and 118 – 210ms, Predictable: 10 – 211ms and 226 – 252ms; all p's < 0.05).

*Topographic analyses.*

Interaction of TACwith contextual factors. In the 5 × 2 × 2 rmANOVA there was a main effect of Map, $F_{(3.1,117.4)}$=11.7, $p$ <0.001, $\eta_p^2$=0.2. Apart from the 3-way Map x Multisensory Relationship × Distractor Onset interaction described in the main text, Map interacted with Multisensory Relationship, $F_{(3.1,117.3)}$=2.8, $p$=0.04, $\eta_p^2$=0.07.

Interaction of MSE with contextual factors. There were three subsequent time-windows where the single-subject data fitting was performed to investigate the interactions between MSE and contextual factors in the ERP topography using the AV-V difference ERPs. The first fitting time-window (35–110ms) showed a main effect of Map, $F_{(1.8,69.7)}$=27.1, $p$ <0.01, $\eta_p^2$=0.4. There was also a nonsignificant trend for a 2-way Map × Distractor Onset interaction, $F_{(1.9,74.6)}$=2.6, $p$ =0.08, $\eta_p^2$=0.06.

The second time-window (110– 190ms) showed a main effect of Map, $F_{(2.5,95)}$=4.2, $p$ <0.01, $\eta_p^2$=0.1. This effect was modulated by both contextual factors separately, aside from the 3-way interaction described in the main text, with a 2-way Map × Multisensory Relationship interaction reliable at $F_{(2.6,99.8)}$=5.5, $p$=0.002, $\eta_p^2$=0.1, and a 2-way Map × Distractor Onset interaction reliable at $F_{(2.8,108.1)}$=3.1, $p$=0.03, $\eta_p^2$=0.08.

Finally, the third time-window (190– 300ms) showed a main effect of Map, $F_{(2.9,109.2)}$=15.2, $p$ <0.001, $\eta_p^2$=0.3. There was also a nonsignificant trend for a 2-way Map × Distractor Onset interaction, $F_{(3.3,126.3)}$=2.4, $p$ =0.07, $\eta_p^2$=0.1.

### Exploratory findings

#### Behavioural results

##### Interaction of TAC x MSE

In the first unexpected finding in this study, the strength of behavioural multisensory enhancement depended on the colour of the distractor, with a 2-way Distractor Colour × Distractor Modality interaction, $F_{(1, 38)}$=6.4, $p$=0.015, $\eta_p^2$=0.2. Capture effects elicited by TCC distractors were larger on AV (46ms) than V trials (38ms) ($t_{(38)}$=4.3, $p$<0.001), while NCC distractors showed no similar effect ($t_{(38)}$<1; see Supplementary Figure 5). However, none of the Tasks showed this effect separately, which suggests that this unexpected finding was driven by a large number of 'participants' (4 Tasks x 39 participants overall = 156). We report the results of the Task-wise analyses below.

*TAC x MSE Interaction within specific Tasks.* We followed up the 2-way interaction of Distractor Colour x Distractor Modality (see Supplemental Figure 5), by analysing it separately within each Task. In Task 1, there was a main effect of Distractor Colour, $F_{(1, 38)}$ = 169.1, $p$ < 0.001, $\eta_p^2$ = 0.8. Post-hoc $t$-tests showed the classical task-set contingent capture effect, whereby TCC stimuli elicited significantly larger spatial cueing effects than NCC stimuli (49ms vs. -1ms, $t$ = 8.9, $p$ < 0.001). The main effect of Distractor Modality was at the level of a nonsignificant trend ($F_{(1, 38)}$ = 3.4, $p$ = 0.07, $\eta_p^2$ = 0.1). There was no significant interaction between these two factors, $F_{(1, 38)}$ = 2.7, $p$ = 0.11, $\eta_p^2$ = 0.07.

In Task 2, there was also a significant main effect of Distractor Colour, $F_{(1, 38)}$ = 141.9, $p$ < 0.001, $\eta_p^2$ = 0.8, where TCCs again elicited larger spatial cueing than NCCs (40ms vs. -3ms, $t$ = 6.6, $p$ < 0.001). However, there was no significant main effect of Distractor Modality, $F_{(1, 38)}$ = 0.5, $p$ = 0.5, $\eta_p^2$ = 0.01, as well as no significant interaction between Distractor Colour and Distractor Modality, $F_{(1, 38)}$ = 2.6, $p$ = 0.11, $\eta_p^2$ = 0.05.

In Task 3, however, both a main effect of Distractor Colour, $F_{(1, 38)}$ = 148.3, $p$ < 0.001, $\eta_p^2$ = 0.8, and a main effect of Distractor Modality, $F_{(1, 38)}$ = 16.8, $p$ < 0.001, $\eta_p^2$ = 0.3 were observed. Post-hoc t-tests revealed that TCC stimuli elicited significantly larger spatial cueing effects than NCC stimuli (38ms vs. -2ms, $t$ = 6, $p$ < 0.001), as did AV stimuli over V stimuli (22ms vs. 13ms, $t$ = 4.2, $p$ < 0.001), replicating the task-set contingent capture and multisensory enhancement effects. Again, there was no significant interaction between these two factors, $F_{(1, 38)}$ = 1.2, $p$ = 0.29, $\eta_p^2$ = 0.03.

Finally, in Task 4, a significant main effect of Distractor Colour, $F_{(1, 38)}$ = 139, $p$ < 0.001, $\eta_p^2$ = 0.8 was observed, with larger spatial cueing for TCCs than NCCs (40.41ms vs. 1.99ms, $t$ = 6.3, $p$ < 0.001). However, there was no significant main effect of Distractor Modality, $F_{(1, 38)}$ = 1, $p$ = 0.3, $\eta_p^2$ = 0.03, as well as no significant interaction between Distractor Colour and Distractor Modality, $F_{(1, 38)}$ = 1.7, $p$ = 0.2, $\eta_p^2$ = 0.06.

##### Interaction of contextual effects with spatial cueing effects

In this study we were interested in the interactions between visual (goal-based; measured by TAC) and multisensory (bottom-up; measured by MSE) attentional control mechanisms interacting with contextual factors related to distractors eliciting the former two traditional effects. While we expected that contextual influences would be visible through modulations of TAC and MSE, de facto, we found behavioural capture effects to be also jointly modulated by the two contextual factors independently of TAC and MSE. That is, we found a two-way interaction between Multisensory Relationship and Distractor Onset, $F_{(1, 38)}$=8.3, $p$=0.007, $\eta_p^2$=0.2.

We first conducted a follow-up analysis as a function of Multisensory Relationship. Post-hoc tests revealed that, for Arbitrary distractors, behavioural capture effects were larger for Unpredictable distractors (24ms) compared to Predictable distractors (19ms), $t_{(38)}$ = 2.8, $p$ = 0.03 (Supplemental

Figure 6). In contrast, capture effects elicited by Congruent distractors were not reliably different as a function of the predictability of distractor onset ($t_{(38)} = 1.7$, $p = 0.1$). The follow-up analysis as a function of Distractor Onset revealed that, for Unpredictable distractors, behavioural capture effects were larger for Arbitrary distractors (24ms) than for Congruent distractors (19ms), $t_{(38)} = 2.7$, $p = 0.006$. Meanwhile, for Predictable distractors, behavioural capture effects were not significantly different between Arbitrary distractors (18ms) and Congruent distractors (21ms), $t_{(38)} = 1$, $p = 0.22$. Thus, the two contextual factors influenced attentional capture directly, such that modulations by Distractor Onset were visible in the absence of a semantic relationship between the visual and auditory characteristics of the distractors, and modulations by Multisensory Relationship were visible when distractor onset could not be predicted.

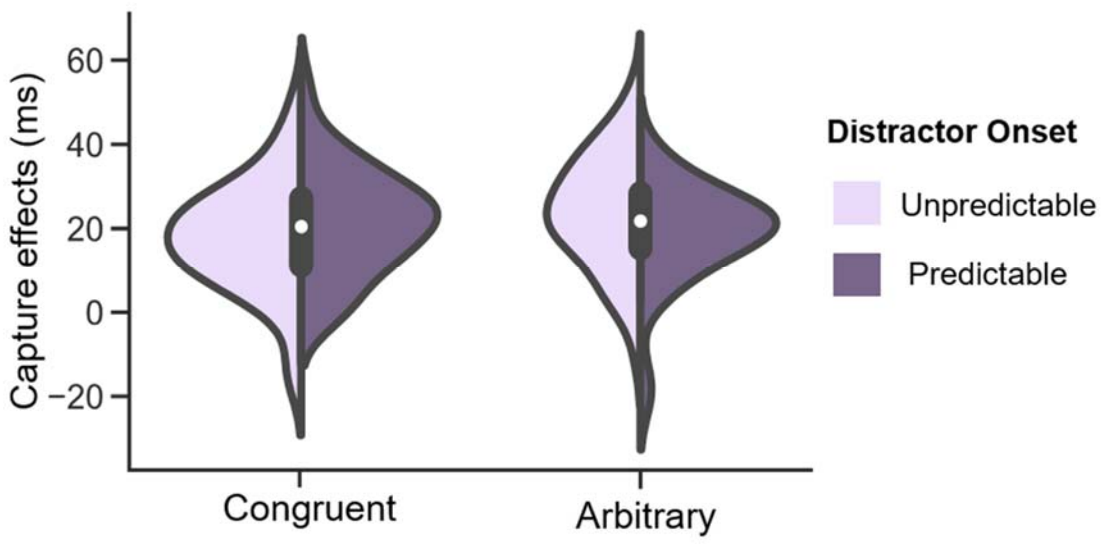

**Supplemental Figure 6.** For Arbitrary distractors, behavioural capture was larger in contexts where Distractor Onset was Unpredictable than Predictable. No such effect was observed for Congruent distractors.

#### ERP analyses

##### The role of lateralised (N2pc-like) brain mechanisms

*Lateralised Electrical Neuroimaging analyses of the N2pc component.* In lateralised ERP topography, map presence over the N2pc (180-300ms) time-window was modulated by contextual factors. Specifically, there was no evidence for a 2-way Map × Multisensory Relationship interaction ($F_{(1.5, 55)} < 0.01$, $p = 1$). However, there was a 2-way Map × Distractor Onset interaction, $F_{(1.8, 67.7)} = 5.9$, $p = 0.006$, $\eta_p^2 = 0.1$. A follow-up analysis of the 2-way Map × Distractor Onset interaction as a function of Distractor Onset revealed that Map2 was present longer for Unpredictable distractors (22ms) than for Predictable distractors (17ms), $t_{(38)} = 2.1$, $p = 0.01$, while Map3 was present longer for Predictable distractors (60ms) than for Unpredictable distractors (52ms), $t_{(38)} = 3.4$, $p = 0.006$. Finally, the 2-way Map × Distractor Onset interaction was further modulated by Multisensory Relationship, $F_{(1.3, 49)} = 5.4$, $p = 0.02$, $\eta_p^2 = 0.1$, the latter interaction mirroring the behavioural results of the two contextual factors interacting directly. The 3-way interaction was first followed up as a function of Multisensory Relationship. Here, for Arbitrary distractors, Map1 was present longer for Unpredictable (53ms) than for Predictable distractors (42ms), $t_{(38)} = 2.4$, $p = 0.007$, and Map3 was present longer for Predictable (63ms) than for Unpredictable distractors (49ms), $t_{(38)} = 3.5$, $p = 0.009$. Meanwhile, for Congruent distractors, it was Map2 that was present longer for Unpredictable (24ms) than for Predictable distractors (14ms), $t_{(38)} = 2.6$, $p = 0.001$. There was likewise a 4-way Multisensory Relationship × Distractor Onset × Distractor Colour × Map interaction observed, but only at a level of a nonsignificant trend, $F_{(1.5, 55.8)} = 3$, $p = 0.06$, $\eta_p^2 = 0.1$. No other main effects ($F$'s < 1) or interactions in the lateralised ERP topography reached statistical significance (all $F$'s < 2, $p$'s > 0.1).

##### The role of non-lateralised brain mechanisms

*Strength-based brain mechanisms.* In GFP we found that Multisensory Relationship and Distractor Onset interacted independently of Target Difference, and did so at 234–249ms. We first followed up the interaction as a function of Multisensory Relationship. For Arbitrary distractors, larger Global field power amplitudes were observed for Predictable than Unpredictable distractors over the tested time-period ($p < 0.05$). For Congruent distractors, GFP was comparable between the levels of Distractor Onset.

#### Supplemental Discussion: Replications

While this was not the main aim of the present study, it is notable that we observed task-set contingent attentional capture (TAC) despite the addition of both the multisensory manipulation (as in Matusz & Eimer 2011) and the two contextual factors to the paradigm. As in traditional literature, target-matching colour (TCC) distractors triggered reliable behavioural capture effects and enhanced mean N2pc amplitudes compared to nontarget matching colour (NCC) distractors (Eimer et al., 2009, Eimer & Kiss, 2008, Kiss et al., 2008a; 2008b). This suggests that this form of visual top-down attentional control of object attentional selection, based on target features (like colour), is an effective, robust form of attentional control, interacting with both multisensory and contextual mechanisms.

Our present results likewise constitute an important replication of the multisensory enhancement of attentional capture (MSE) effect demonstrated by Matusz and Eimer (2011). Given the strong feature-specific visual top-down attentional demands of the experimental task, the evidence of behavioural multisensory enhancement of visual attentional capture supports the idea put forward by Matusz and Eimer (2011) that these modulations of visual attentional selection can occur involuntarily and independently of task demands (see also Matusz et al. 2015). We did indeed observe again the overall multisensory enhancement of attentional capture (increase of attentional capture for colour distractors on tone-present trials), reflected by a main effect of Distractor Modality. However, unlike the original study (Matusz & Eimer 2011), now the multisensory enhancement was apparently dependent on the colour matching between target and the distractor, as shown by a significant 2-way Distractor Colour x Distractor Modality interaction. More specifically, multisensory enhancement of behavioural attentional capture was found for target-matching (TCC) distractors, but not for nontarget-matching (NCC) distractors. However, when we probed the Distractor Colour x Distractor Modality interaction within each of the Tasks, the 2-way Distractor Colour x Distractor Modality interaction was no longer present. It was not present even in Task 2 that was the closest to the original study, in that distractors were Predictable and Arbitrary. While care is needed when interpreting such null results, one possibility is that, in fact, in the real world, bottom-up multisensory and top-down goal-directed attentional control processes interact (c.f. Desimone & Duncan (1995), for qualification of 'bottom-up' processes as potentially involving feedback signals). Perhaps goal-based attentional control processes do benefit from increased salience of non-semantically related multisensory objects. However, it is also possible that such interactions are subtle and/or the current paradigm is not optimal for gauging them.

Furthermore, the results from electrical neuroimaging analyses of the non-lateralised responses provide a novel insight into the attentional selection of audio-visual objects. Specifically, even when such objects are irrelevant to the task, multisensory relationships modulate their processing, they do so from earlier perceptual stages to later cognitive stages, and through interactions with other contextual factors, such as stimulus (temporal predictability.
